# Detailing organelle division and segregation in *Plasmodium falciparum*

**DOI:** 10.1101/2024.01.30.577899

**Authors:** Julie M.J. Verhoef, Cas Boshoven, Felix Evers, Laura J. Akkerman, Barend C.A. Gijsbrechts, Marga van de Vegte-Bolmer, Geert-Jan van Gemert, Akhil B. Vaidya, Taco W.A. Kooij

## Abstract

The malaria causing parasite, *Plasmodium falciparum*, replicates through a tightly orchestrated process termed schizogony, where approximately 32 daughter parasites are formed in a single infected red blood cell and thousands of daughter cells in mosquito or liver stages. One-per-cell organelles, such as the mitochondrion and apicoplast, need to be properly divided and segregated to ensure a complete set of organelles per daughter parasites. Although this is highly essential, details about the processes and mechanisms involved remain unknown. We developed a new reporter parasite line that allows visualization of the mitochondrion in blood and mosquito stages. Using high-resolution 3D-imaging, we found that the mitochondrion orients in a cartwheel structure, prior to stepwise, non-geometric division during the last stage of schizogony. Analysis of focused ion beam scanning electron microscopy (FIB-SEM) data confirmed these mitochondrial division stages. Furthermore, these data allowed us to elucidate apicoplast division steps, highlighted its close association with the mitochondrion, and showed putative roles of the centriolar plaques (CPs) in apicoplast segregation. These observations form the foundation for a new detailed mechanistic model of mitochondrial and apicoplast division and segregation during *P. falciparum* schizogony and pave the way for future studies into the proteins and protein complexes involved in organelle division and segregation.

## Introduction

Malaria is a devastating parasitic disease causing an estimated 249 million cases resulting in approximately 608,000 deaths in 2022, especially in children under 5 years old^1^. *Plasmodium falciparum* is the most virulent parasite species causing malaria. Continued emergence of resistant parasites to antimalarial drugs is a major problem for global malaria control and necessitates continued development of novel antimalarials.

The malaria parasite harbors a unique mitochondrion that differs greatly from the human mitochondrion at a molecular and functional level^2^. While the most prominent role of the mitochondrion in humans is respiration and consequent energy conversion, in the disease-causing asexual blood stages of *P. falciparum* the respiratory chain appears to be exclusively essential to support pyrimidine biosynthesis^3^. It is only during preparation for transition to the mosquito vector where sexual reproduction takes place, that canonical mitochondrial functions such as the tricarboxylic acid cycle (TCA) cycle and the oxidative phosphorylation (OXPHOS) pathway become more abundant and critical^4,5^. Because of these differences, it is not surprising that this organelle is the drug target of several anti-malarial compounds, such as atovaquone, DSM265, proguanil and ELQ300^6,7^.

Host and stage transitions are commonplace in the complicated life cycle of *Plasmodium* parasites. During erythrocytic asexual replication, one parasite is segmented into approximately 32 merozoites through a tightly orchestrated process called schizogony. Cell division happens on a much larger scale in mosquito and liver stages, where one parasite is divided into thousands or even tens of thousands of daughter parasites. During *P. falciparum* cell division, the single parasite mitochondrion needs to be properly divided and distributed among the daughter cells^9^. During parasite development in asexual blood stages, the tubular mitochondrion elongates and forms a large, branched network that stretches throughout the parasite^10^. Only during the final stages of schizogony, once nuclear division is completed, does the mitochondrion undergo rapid fission^11^. The apicoplast, another essential single copy organelle of secondary endosymbiotic origin, forms a comparable branched network, but divides prior to mitochondrial fission during blood-and liver-stage replication^10,12^. To produce viable offspring, the parasite has to ensure that each daughter parasite has a complete set of these organelles. However, so far a detailed view of these processes and the mechanisms involved is lacking.

We aimed to capture the process of mitochondrial division in these multinucleated cells in detail using different imaging methods. However, this comes with several challenges. Firstly, imaging the small-sized parasites (1-7 μm diameter), and the even smaller organelles within the parasites, requires the use of super-resolution imaging techniques. Secondly, visualization of the mitochondrion requires a specific fluorescent marker or dye. Mitochondrial dyes, such as Rhodamine123 and MitoTracker^TM^, have been widely used in the field^13^. These dyes rely on membrane potential to enter the mitochondrion and are therefore also used as a viability marker^14^. However, eight of these dyes were tested in a drug screen all showing IC50 values below 1μM with three, Mito Red, DiOC_6_, and Rhodamine B being highly active against *P. falciparum* with IC50 values below 30 nM^15,16^. Additionally, in our hands MitoTracker signal can be diffuse, and therefore limit the resolution that is needed for the visualization of mitochondrial fission. Hence, we aimed to develop a reporter parasite line which harbors a fluorescent mitochondrial marker that allows imaging of this organelle in live and fixed conditions in all life-cycle stages of *P. falciparum*. To do this, we deployed a similar strategy that has been used successfully in the rodent model *Plasmodium berghei*^17,18^. The targeting signal of the known mitochondrial protein HSP70-3 was fused with a fluorescent protein and integrated in a silent intergenic locus (SIL)^19^. Expression of this mitochondrial-localized fluorescent protein allowed visualization of the organelle during imaging of asexual, sexual and mosquito stages. Using high resolution confocal microscopy, we were able to make a detailed 3D map of different mitochondrial fission stages during schizogony in asexual blood stages. Focused ion beam scanning electron microscopy (FIB-SEM) image stacks from Evers *et al.* were used to confirm these mitochondrial fission stages with high detail^20^. This also allowed us to study apicoplast division and highlighted the potential role of the centriolar plaques (CPs) in apicoplast segregation. These different microscopic approaches empowered us to put forward a detailed model for mitochondrial and apicoplast division and distribution during the final stages of schizogony.

## Results

To acquire a detailed understanding of mitochondrial fission and distribution, we set out to capture this process throughout the *Plasmodium* life cycle by combining different microscopy approaches. We stained mature blood-stage wild-type *P. falciparum* NF54 strain parasites with two different MitoTracker dyes and used these to visualize the mitochondrion in fixed confocal imaging. Surprisingly, both MitoTracker dyes showed a discontinuous, punctate staining pattern (Figure 1A). FIB-SEM studies have confirmed the prevailing notion that the mitochondrion is a single, branched network during these schizont stages^20^. While this observation may arise from crosslinking of the MitoTracker dyes to specific proteins and aggregations thereof resulting from the fixation process, we concluded that the punctate staining pattern is likely an artifact and consequently limits our ability to dissect and visualize the process mitochondrial fission. To address this, we developed a new fluorescent mitochondrial marker that can be used for imaging live and fixed samples (Figure S1).

**Figure 1.**
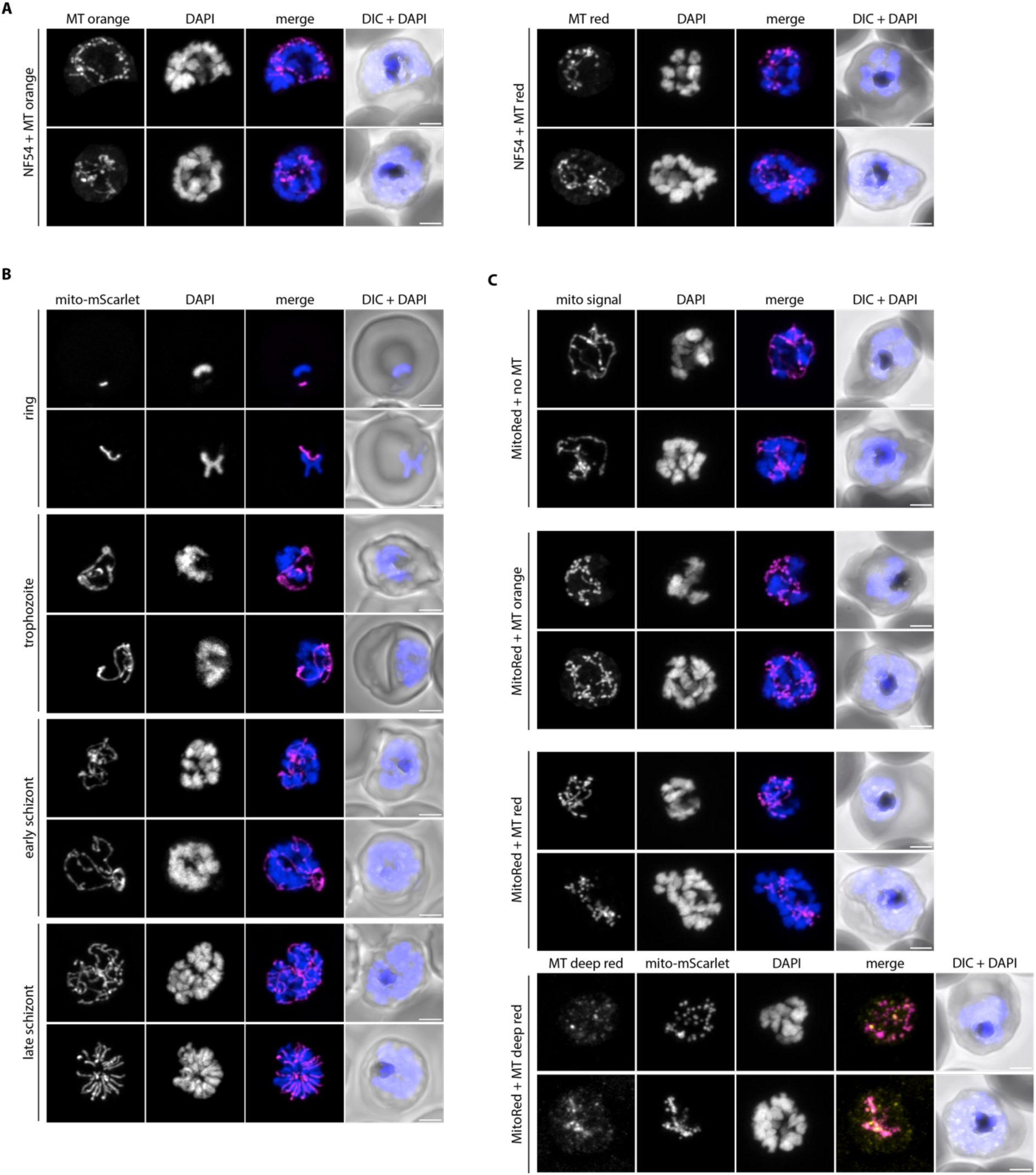
Comparison of MitoTracker and a new mitochondrial marker for fluorescence imaging. A) Fluorescent imaging of WT parasites stained with MitoTracker Orange CMTMRos (MT orange) or MitoTracker Red CMXRos (MT red). B) Fluorescence microscopy of MitoRed. The mito-mScarlet signal was observed in all asexual life-cycle stages including rings, trophozoites, early and late schizonts. No antibody staining was used and fluorescent signal observed is exclusively the mito-mScarlet signal. C) Fluorescence microscopy of MitoRed, either unstained (no MT) or stained with MT orange, MT red or MitoTracker Deep Red FM (MT deep red). Mito signal is the combined MitoTracker and mito-mScarlet signal that is observed in this channel. DAPI (blue) is used to visualize DNA and DIC (differential interference contrast) for general cellular context. All images are maximum intensity projections of Z-stacks (41 slices, 150 nm interval) taken with Airyscan confocal microscope. Scale bars, 2 μm.

### Design and generation of a new mitochondrial marker parasite line

We designed a mitochondrial marker that consists of the promotor and mitochondrial targeting sequence of the gene encoding the mitochondrial heat shock protein 70 (HSP70-3, PF3D7_1134000), fused to an mScarlet red fluorescent protein (Figure S1A). HSP70-3 was selected based on its high and consistent expression profile throughout the whole life cycle and has been successfully used for the same purpose in *P. berghei*^17,18^. We aimed to stably integrate this fluorescent marker in the *P. falciparum* genome, without affecting any normal biological processes and parasite growth throughout the parasite life cycle. Selection of the new integration site, SIL7, is described extensively in Supplemental Information S1. The integration plasmid was transfected into NF54 parasites together with two different Cas9 guide plasmids directed at the SIL7 site. Successful integration of the mitochondrial marker and absence of WT parasite contaminations were confirmed by integration PCR (Figure S1B). A growth assay showed no difference in growth of the mitochondrial reporter line, MitoRed, compared to WT parasites in asexual blood stages (Figure S1C).

### Characterization of the MitoRed parasite line

To visualize the mitochondrial marker, asexual blood-stage MitoRed parasites were fixed and used for fluorescent imaging. The fluorescent signal was well preserved after fixation and no antibody staining was required for mitochondrial visualization in all asexual blood stages (Figure 1B). To assess whether the punctate mitochondrial morphology observed after MitoTracker staining was an imaging artifact or a morphological aberration caused by the dye, we stained MitoRed parasites with three different MitoTracker dyes. Discontinuous, punctate mitochondria were observed in all MitoRed parasites stained with MitoTracker, while this was not observed in unstained MitoRed parasites (Figure 1C). The effect was less pronounced during live imaging of MitoTracker stained parasites (Figure S2). While there is an obvious imaging artifact following fixation of MitoTracker-stained blood-stage *P. falciparum* parasites, the altered MitoRed signal in the presence of the dye might even suggest possible changes in mitochondrial morphology.

### Mitochondrial dynamics during gametocyte development and activation

To study mitochondrial dynamics throughout the malaria parasite life cycle, MitoRed parasites were induced to form gametocytes, which were fixed for microscopy on day 5, 7, 10, and 13 post induction. Parasites were stained for α tubulin to distinguish male and female gametocytes in stage IV and V. For each stage, between 11-19 parasites were imaged over two independent experiments and described observations were consistent over all analyzed parasites. In stage II and III gametocytes, the mitochondrion appears as a small knot that increases slightly in size when gametocytes become more mature (Figure 2A). This is consistent with our FIB-SEM data^20^. Evers *et al*. also showed that gametocytes have multiple mitochondria already from early gametocyte development onwards. Although light microscopy does not provide the resolution or ability to show membrane boundaries to distinguish the multiple mitochondria in stage II and III gametocytes, in stage IV gametocytes we could clearly observe separate mitochondria in both male and female gametocytes (Figure 2A, Figure S3A). There is no clear difference in stage IV gametocytes between male and female mitochondria. However, in stage V gametocytes the mitochondria in males appear slightly more dispersed, while the female mitochondria remain compact (Figure 2A, Figure S3B). Data from Evers *et al.* support this and showed consistently smaller volume and more loosely packed mitochondria in males compared to females^20^. When gametocytes are taken up by the mosquito via a blood meal, they are activated and transform into extracellular male and female gametes. While the female gametocyte develops into a single macrogamete, male gametocytes form up to eight flagellated microgametes. This transformation is triggered by a temperature drop, an increase in pH, and xanthurenic acid present in the mosquito midgut^21^. Upon *in vitro* activation, the difference between male and female mitochondria becomes more evident. Mitochondria in females remain in a compact knot while the parasite rounds up (Figure 2B). Interestingly, in males the mitochondria become smaller and more dispersed, and sometimes round up to small bean-like structures (Figure 2B, Figure 2D). This process already starts 2 min after activation. While this particular activation experiment was performed on a gametocyte culture that did not exflagellate for unclear reasons, it was repeated twice, and very similar results were found in exflagellating males (n=19) (Figure 2C). There was no significant difference between number of exflagellation events in MitoRed parasites compared to NF54 parasites (Figure S4A). An association of mitochondria with flagella is not uncommon and can also be observed in *e.g.* kinetoplastids, such as *Trypanosoma* and *Leishmania spp*, where the mitochondrion resides at the base of the flagellum, and in human sperm cells, where the mitochondrion wraps around the base of the flagellum to provide energy for flagellar movement^22^. We found close apposition of the dispersed mitochondria to the axonemal tubulin in all 19 exflagellating males that were analyzed (Figure S4B, S4C).

**Figure 2.**
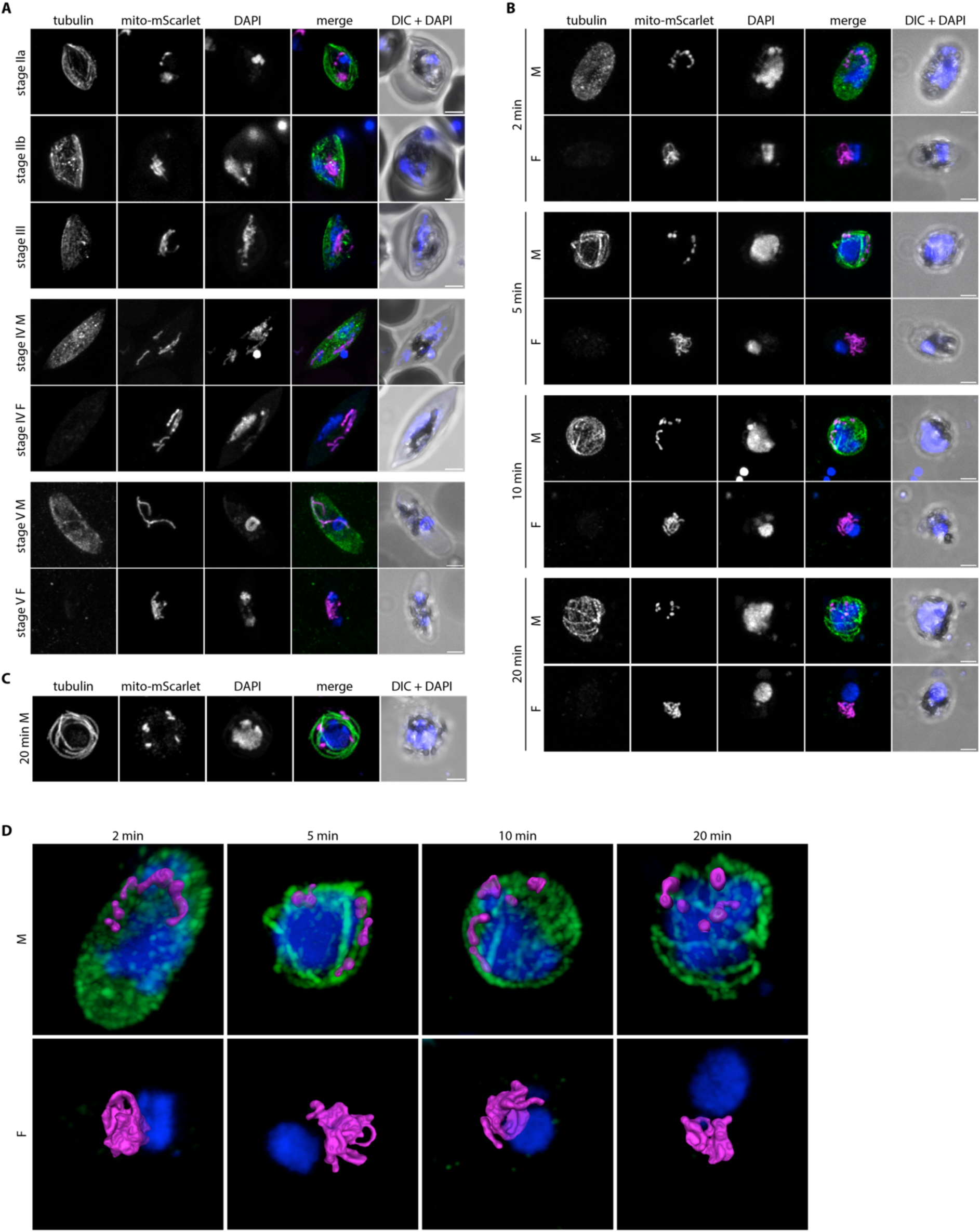
Mitochondrial dynamics during gametocyte development and activation. A) Immunofluorescence assay on MitoRed gametocytes stages IIa, IIb, III, IV, and V, stained with anti-β-tubulin (green) and DAPI (DNA, blue). The mito-mScarlet signal is shown in magenta. In stage IV and V, male (M) and female (F) gametocytes are distinguished based on the intensity of the tubulin signal (males high, females low). B) Immunofluorescence assay on MitoRed parasites during different stages of gametocyte activation (2, 5, 10 and 20 min after activation). C) Immunofluorescence assay on MitoRed exflagellating male gamete 20 min after activation. A-C) Images are maximum intensity projections of Z-stacks (41 slices, 150 nm interval) taken with an Airyscan confocal microscope. Scale bars, 2 μm. D) 3D visualization of male and female MitoRed parasites 2, 5, 10, and 20 min after activation. The mito-mScarlet fluorescent signal is segmented based on manual thresholding.

### Mitochondrial dynamics in mosquito stages

In the mosquito midgut, the male microgamete seeks out a female gamete for fertilization. After fertilization, the zygote takes one day to transform into a motile ookinete, which can traverse the midgut epithelium and differentiate into an oocyst. This oocyst expands and motile sporozoites are formed within the oocyst. When fully matured, the oocyst will burst and sporozoites will egress, spread through the hemolymph system, and invade the mosquito salivary glands. During oocyst development, the parasite mitochondrion has to expand enormously and then be divided over thousands of daughter sporozoites. However, very little is known about mitochondrial dynamics and only few studies have visualized the mitochondrion during these stages^17,23,24^.

To explore if MitoRed parasites develop normally in the mosquito and to visualize mitochondrial morphology, mature MitoRed gametocytes were fed to *Anopheles stephensi* mosquitoes. One day after the feed, the mosquito blood bolus was extracted and stained with anti-*Pf*s25 conjugated antibodies to visualize ookinetes by live microscopy. We distinguished different stages of ookinete maturation as described by Siciliano *et al.*^25^. Due to the resolution limit of light microscopy, it was difficult to tell if there were one or multiple mitochondria as observed in gametocyte stages. Since we did not find evidence for the presence of multiple mitochondria in these ookinete development stages, we will refer to it as “the mitochondrion” in the coming paragraph, although we cannot rule out the presence of multiple mitochondria. During earlier stages of ookinete development (II), when ookinetes have a short protuberance attached to the round body, the mitochondrion resides in the round body (Figure 3A, 3B). When the protuberance starts to elongate further, one elongated mitochondrial branch stretches out and reaches into the protuberance. In stage III ookinetes, the mitochondrion stretches out further into the growing protuberance, spiraling out from the round body. We could not find clear stage IV ookinetes where the protuberance is at its full length, which could be explained by the swift development from stage IV to V ookinetes as was observed by Siciliano *et al.*^25^. However, in the mature stage V ookinetes, the mitochondrion appears as a tight knot in the main parasite body.

**Figure 3.**
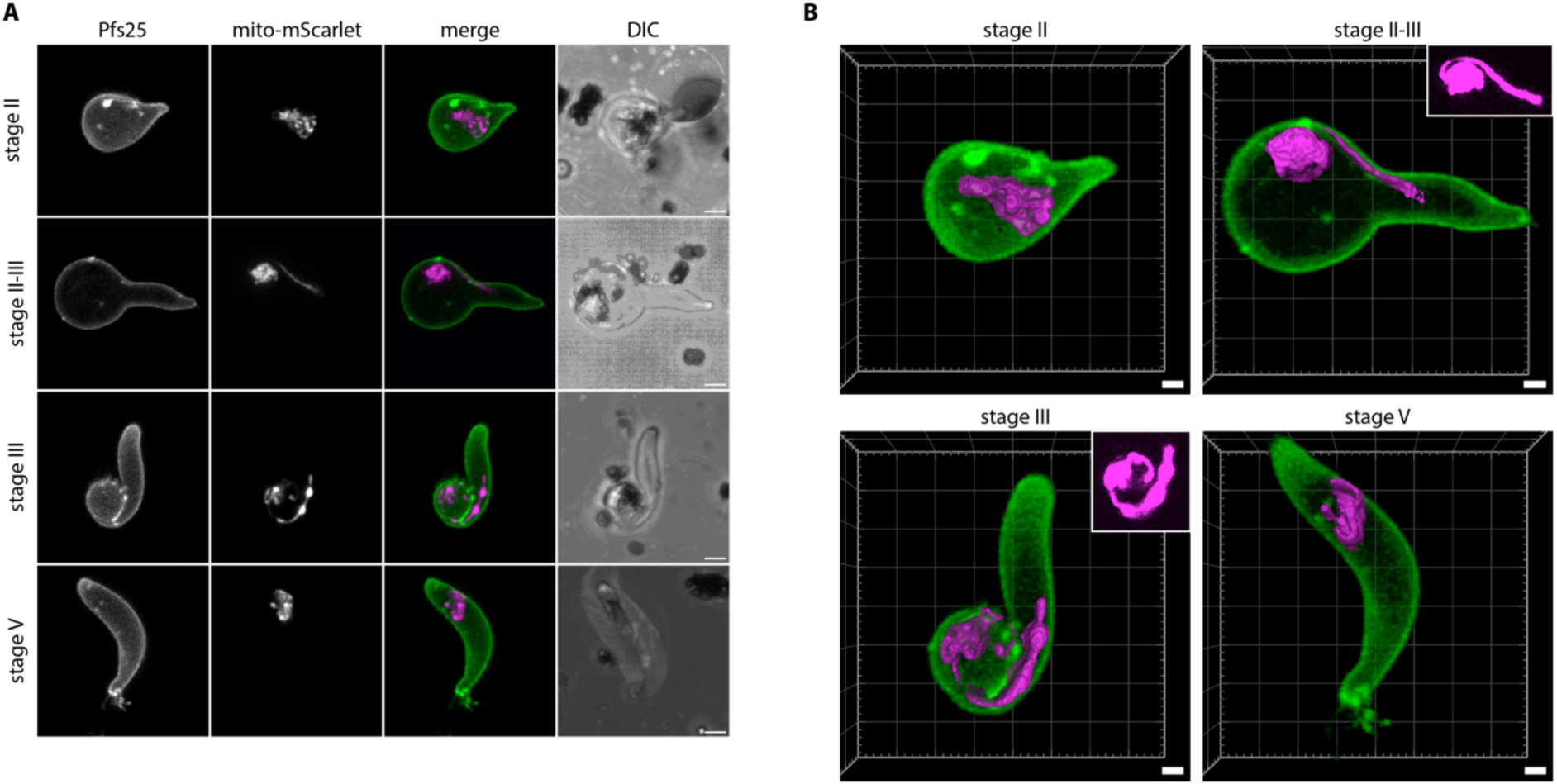
Mitochondrial dynamics during ookinete development. A) Live imaging of MitoRed ookinetes one day after mosquito feed. Different stages of ookinete maturation (II – V) were distinguished based on description by Siciliano et al.^25^. Cells were stained with an Alexa fluor 488 conjugated anti-Pfs25 antibody to visualize parasite outline (green). Images are maximum intensity projections of Z-stacks (30 slices, 185 nm interval) taken with an Airyscan confocal microscope. Scale bars, 2 μm. B) 3D visualization of different ookinete maturation stages. The mito-mScarlet fluorescent signal is segmented based on manual thresholding. Two smaller images in upper right corner of stage II-III and stage III are crops of the mitochondrial fluorescent signal with increased brightness and contrast. Scale bars, 1 μm.

At day 7, 10, and 13 after infection, mosquitoes from a feed with an infection rate of 100% and an average of 5 oocysts/mosquito were dissected and midguts were used for live confocal microscopy. At day 7, small oocysts (∼10 μm diameter) were observed with a branched mitochondrial network stretched out throughout the cell (Figure S5A). Segmentation of the fluorescent signal based on manual thresholding indicated that the mitochondrion consisted of one continuous structure. Day 10 oocysts were much larger (∼35 μm diameter) and the mitochondrial mesh-like network appeared more organized, also localizing to areas directly below the oocyst wall (Figure S5B). At day 13, oocysts of various sizes were observed. Some large oocysts (∼70 μm diameter) showed a highly organized mitochondrial network, where mitochondrial branches were organized in a radial fashion around a central organizational point (Figure S5C). We named these points mitochondrial organization centers (MOCs). At least tens of these MOCs could be observed per cell. Some smaller oocysts (∼35 μm diameter) at day 13 showed structures that looked like beginning MOCs (Figure S5D). However, several small oocysts showed a dispersed, globular mitochondrial signal, which we interpreted as unhealthy or dying parasite (Figure S5E). While several free sporozoites were observed in dissected midguts and salivary glands on day 16 (data not shown), we never observed an oocyst containing fully mature sporozoites with a divided mitochondrion or an infected salivary gland on day 16 and 21 after infection. This indicates that sporozoites are produced and released into the hemocoel, however, they have a health defect that prevents them from infecting the salivary glands. Possibly the mitochondrial marker or the integration in the SIL7 locus causes issues for sporozoite development. We conclude that the MitoRed line is a great tool for mitochondrial visualization in asexual blood stages, gametocytes stages, and mosquito stages up until late oocysts (Supplemental Information S1) but that for studies later in the life cycle other tools need to be developed and tested.

### Mitochondrial division during schizogony in asexual blood stages

Next, we aimed to use MitoRed for live visualization of mitochondrial division during schizogony in asexual blood stages. The biggest advantage of live imaging is that one parasite can be followed over time to capture mitochondrial fission events chronologically. Unfortunately, this proved to be challenging. All parasites imaged in several experiments for a duration exceeding 60 min exhibited significant morphological alterations, including mitochondrial swelling, fragmentation, and formation of vesicle-like structures, which indicate an unhealthy or dying parasite (Figure S6A). Additionally, we frequently observed parasites egressing from their red blood cells (RBCs) after approximately 45 min of imaging, indicating that imaged parasites are unhealthy (Figure S6B). Optimizing imaging conditions by reducing laser power, increasing time interval, better temperature control, and gassing of the imaging chamber with low oxygen mixed gas (3% O_2_, 4% CO_2_), did not improve parasite health during imaging. Therefore, we decided to go for a fixed imaging approach to capture mitochondrial division in asexual blood stages.

To capture mitochondrial fission, MitoRed parasites were tightly synchronized and fixed between 32-36 and 36-40 hours after invasion. In our culture system, MitoRed parasites have a replication cycle of approximately 40 hours, so we captured the last eight hours of schizont maturation before merozoite egress from the RBC. In order to distinguish the precise stage of schizont maturation, we included an anti-GAP45 antibody staining. Glideosome associated protein 45 (GAP45) is an inner membrane complex (IMC) protein and is important for RBC invasion^26,27^. IMC formation starts at the apical end of a developing merozoite during schizogony and continues to develop until it fully encapsulates the daughter merozoite with its own IMC membrane^28,29^. We used the stage of IMC formation and therefore merozoite segmentation as a marker for the maturity and age of the schizonts. Based on IMC and DNA staining, we differentiated four stages of schizont maturation: pre-segmentation (n=6), early-segmentation (n=9), mid-segmentation (n=15), and late-segmentation (n=10, Figure 4).

**Figure 4.**
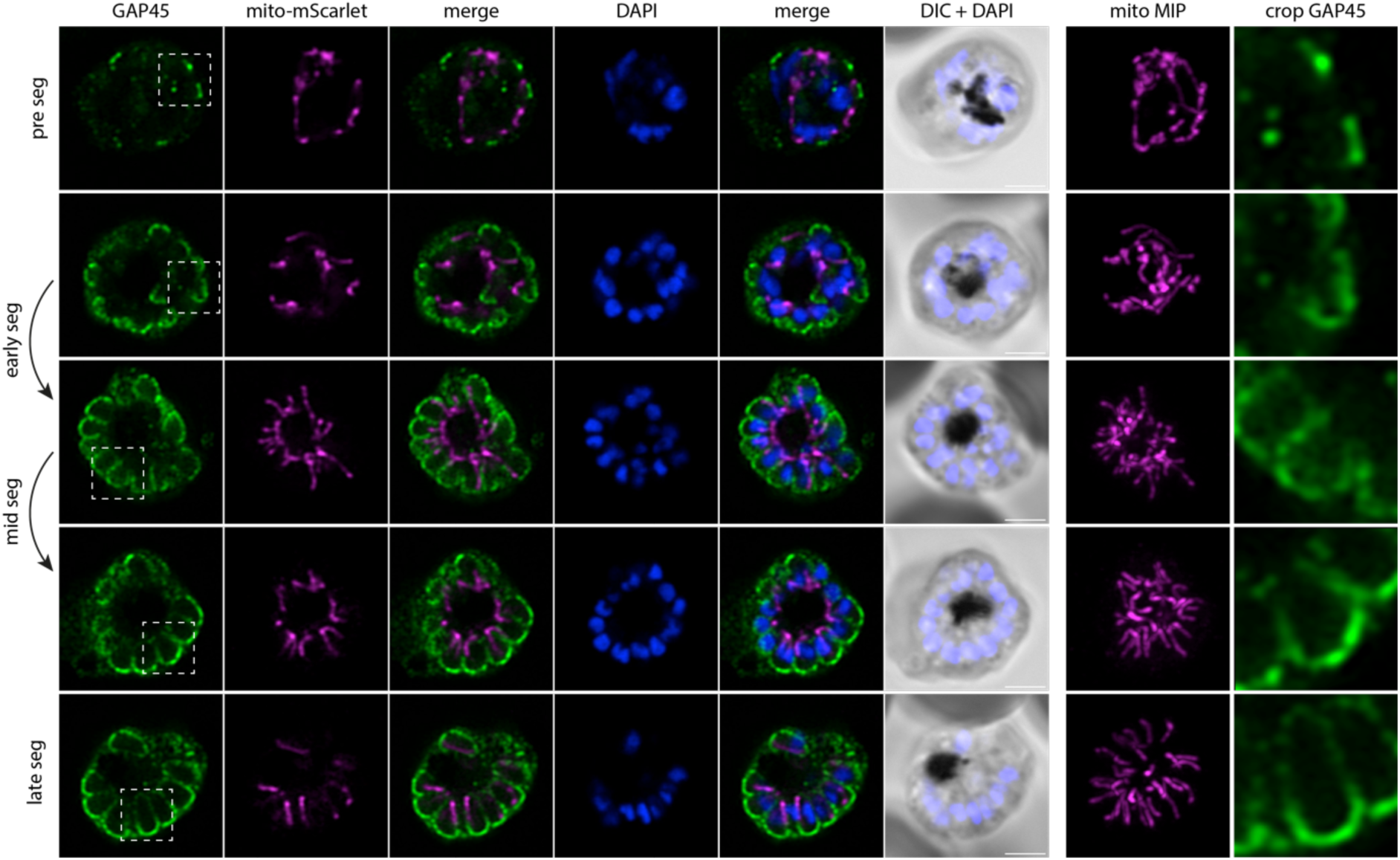
Mitochondrial fission in asexual blood-stage parasites. Immunofluorescence assay on MitoRed schizonts stained with anti-GAP45 antibody (green) to visualize IMC and DAPI (DNA, blue). The mito-mScarlet signal is shown in magenta. Four different stages of schizont maturity are distinguished: <u>pre-segmentation</u> (pre seg), schizonts still undergo nuclear division (nuclei are large and irregularly shaped) and there is no, or very little IMC staining without clear curvature. <u>Early-segmentation</u> (early seg), schizonts have (almost completely) finished nuclear division (nuclei are small and round), there is a clear IMC signal that has a curved shape at the apical end of the forming merozoites but is less than half-way formed. <u>Mid-segmentation</u> (mid seg), the IMC of the segmenting merozoites in these schizonts is more than half-way formed, but there is still a clear opening at the basal end of the merozoite. <u>Late-segmentation</u> (late seg), in these schizonts the IMC seems to be completely formed with no clear opening at the basal end of the forming merozoites. Images are single slices of a Z-stack taken with an Airyscan confocal microscope. Images of the mito-mScarlet signal in the seventh column are maximum intensity projections (MIPs) (41 slices, 150 nm interval). Images in the eighth column are crops of the GAP45 signal depicted in the first column, indicated by the dotted-line areas. Scale bars, 2 μm.

We generated and classified Z-stack images of 40 schizonts, which allowed us to reconstruct a timeline of mitochondrial fission. During pre- and early-segmentation stages, the branched mitochondrial network stretches throughout the parasite. Only at the end of early-segmentation stages, when the IMC is approximately halfway formed, the mitochondrion is oriented around the food vacuole in the center of the parasite with its branches pointing outwards in a radial fashion, creating a “cartwheel”-like structure (Figure 4). As the IMC progresses further and schizonts enter the mid-segmentation stage, this mitochondrial cartwheel structure is divided into smaller fragments, which maintain their radial branch orientation into the segmenting merozoites. Only when IMC formation appears complete, did we observe mitochondria that are entirely divided and distributed over the daughter merozoites. This highlights the extremely late timing of this process. These mitochondrial division stages were confirmed in a second, independent 3D imaging experiment (Figure S7).

To further quantify the numbers and sizes of mitochondria and to create 3D renderings of the mitochondrial network throughout segmentation, we utilized threshold-based masking of the fluorescent signal (Figure 5). During pre- and early-segmentation stages, the mitochondrial network consists of one large fragment (between 7-14 μm^3^), often with 1-3 smaller fragments (<1.5 μm^3^) (Figure 5AI, 5AII). As evident from our FIB-SEM data (Figure S10A), the mitochondrion features constricted regions, characterized by notably reduced diameters. Hence, the smaller fragments observed during these stages are likely not autonomous but caused by the reduced fluorescent marker intensity in the constricted regions. At the end of early-segmentation stages when the IMC is almost halfway formed, the mitochondrial network starts to orient itself in a radial fashion around the center of the parasite (Figure 5AII), consistent with the 2D image analysis (Figure 4). During mid-segmentation stages, the radial mitochondrial branches elongate further into the developing merozoites, and the large mitochondrial fragment is divided in smaller fragments at the center of the cartwheel structure (Figure 5AIII). There is a slight increase in number of mitochondrial fragments per parasite, specifically the “intermediate” sized mitochondrial fragments of 1-4 μm^3^. Only in the last stage of merozoite segmentation, there is a big increase in the number of mitochondrial fragments (Figure 5C). Of note, there appears to be no correlation between this number and the number of nuclei in the parasites (Figure 5B, 5C). A likely technical explanation is the limited Z-resolution of light microscopy and the different nuclear and mitochondrial segmentation methods. When mitochondrial fragments are located closely above each other, the limited Z-resolution in combination with threshold-based masking can cause the adjacent fragments to appear as one continuous structure. Therefore, the number of mitochondrial fragments per schizonts will be underestimated in these late schizont stages. The nuclei on the other hand were segmented through an automated (spherical) object detection algorithm which does not have this problem. Even when the IMC formation appears to be completed based on the GAP45 staining, only 20% of cells appear to have concluded mitochondrial fission as indicated by exclusively containing homogenously small sized mitochondrial fragments (<1.0 μm^3^) (Figure 5AIV). During the late segmentation stage, some parasites still have a large mitochondrial fragment of more than 5 μm^3^, while others only have small and intermediate sized mitochondrial fragments (Figure 5D). This suggest that division of the mitochondrial cartwheel structure into small fragments is a fast, stepwise process that does not happen in a 2^n^ progression and happens only in the final moments of merozoite segmentation.

**Figure 5.**
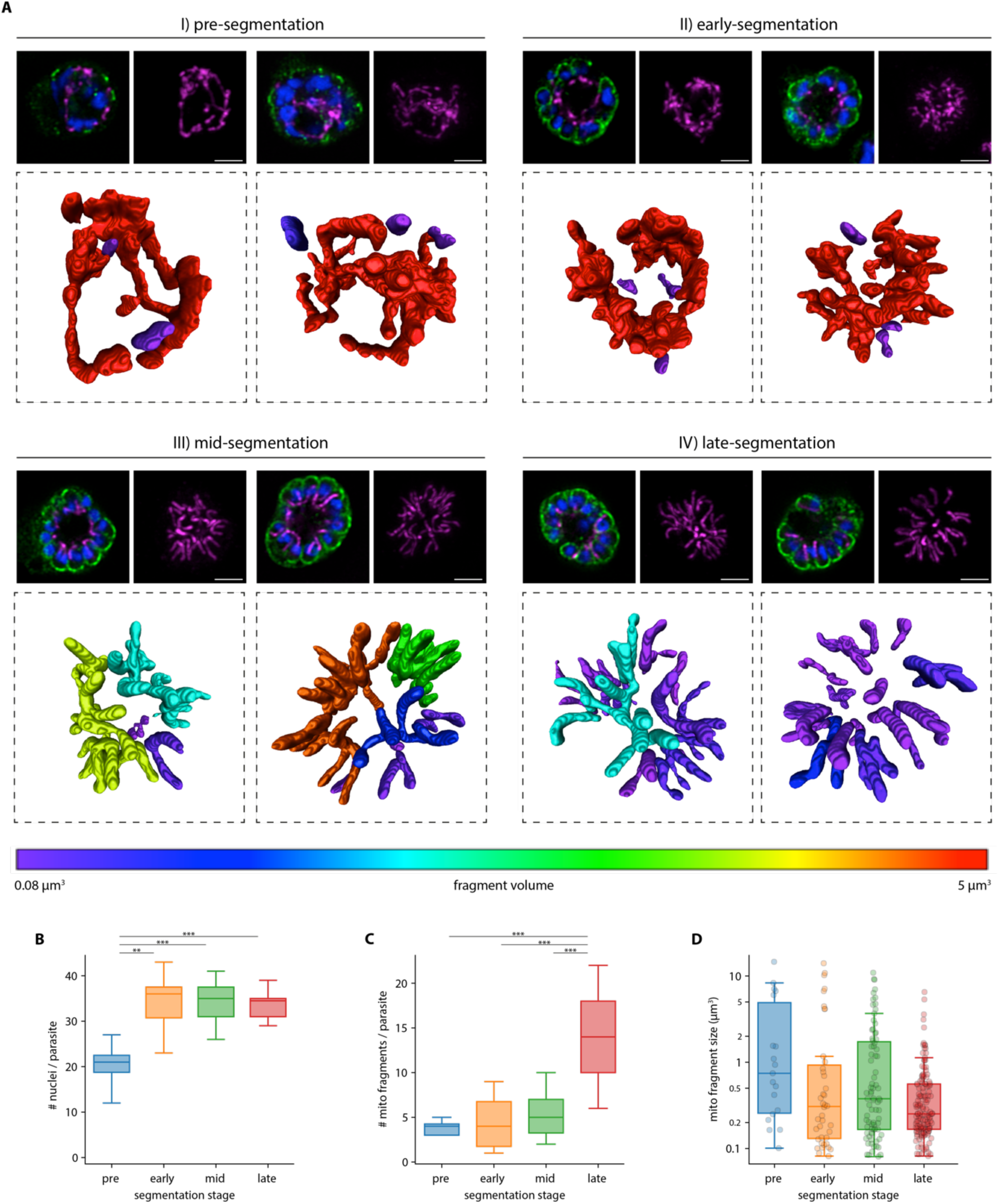
3D analysis of mitochondrial fission stages during schizogony. A) 3D visualization of mitochondrial segmentations based on thresholding of the mito-mScarlet signal in Arivis image analysis software. Smaller images in top row are a single slice of the Z-stack with anti-GAP45 labelling (IMC, green), DAPI (DNA, blue) and mito-mScarlet (magenta), and a maximum intensity projection of the mito-mScarlet signal. The larger bottom picture is a 3D visualization of the segmented mitochondrial signal. The color of the mitochondrial fragment represents the size of this fragment, as is shown in the color bar at the bottom. Two representative parasites are depicted for each of the four segmentation stages defined in Figure 4. Scale bars, 2 μm. B) Boxplot indicating the number of nuclei per parasite in the different segmentation stages. **, p<0.01; ***, p < 0.001. C) Boxplot indicating the number of mitochondrial fragments per parasite in the different segmentation stages. ***, p < 0.001. D) Boxplot indicating the size of the mitochondrial fragments in the different segmentation stages.

### Visualization of mitochondrial and apicoplast division using volume electron microscopy

Although the use of light microscopy allowed us to reconstruct mitochondrial fission in good temporal resolution, its limited spatial resolution and reliance on indirect staining leaves some questions unanswered. Our recent volume electron microscopy study detailed parasite organelle structures at a nanometer resolution bringing many new insights to the light^20^. Here, we reused the underlying FIB-SEM data, which besides gametocytes also contains asexual blood-stage parasites from different stages, to examine mitochondrial fission with high resolution. Asexual parasites in different stages of schizogony were selected and organelles including nuclei, mitochondrion, and apicoplast were segmented for 3D rendering (detailed description per parasite in Tables S2 and S3). The mitochondrion and apicoplast can be recognized by their tubular shape in addition to the double membrane of the mitochondrion and the thicker appearance of the four membranes of the apicoplast. In line with the results from our light microscopy experiments, the mitochondrion is a large, branched network stretched throughout the cell in early schizont stages before segmentation has started (Figure 6, Movie 1). The apicoplast is also a branched network, however, it is much smaller than the mitochondrion (Table S2). The apicoplast network is divided into smaller fragments of different sizes when nuclear division is still ongoing and IMC formation has started (Figure 7, Table S3, Movie 4). When nuclear division is finishing and the IMC envelops part of the nucleus, apicoplast division is completed (Figure 6, Movie 5). The mitochondrion starts to orient its branches in a radial fashion towards the developing merozoites. When nuclear division is completely finished and the IMC envelops most of the nucleus, the mitochondrion forms a clear cartwheel structure with its branches pointing into the developing merozoites (Movie 6). During late segmentation stages, where only a small opening is connecting the merozoite to the residual body, the mitochondrion is divided into smaller fragments of various sizes (Movie 7). While some mitochondrial fragments have a volume comparable with the mitochondria in a fully segmented parasite (0.016 - 0.036 μm^3^), other fragments are still 2-4 times that volume (FigureS8).* These larger mitochondrial fragments have several branches that are pointing into developing merozoites but are still connected to each other outside the merozoites. In an almost fully segmented schizont, where most merozoites are fully developed and only few merozoites are still connected to the residual body through a small opening, the mitochondrial division is completed and the number of mitochondrial fragments is the same as the number of merozoites (Movie 8). These findings corroborate our light microscopy data and confirm the mitochondrial division stages, position of relevant structures not stained in light microscopy, and timing of mitochondrial and apicoplast division during schizogony.

**Figure 6.**
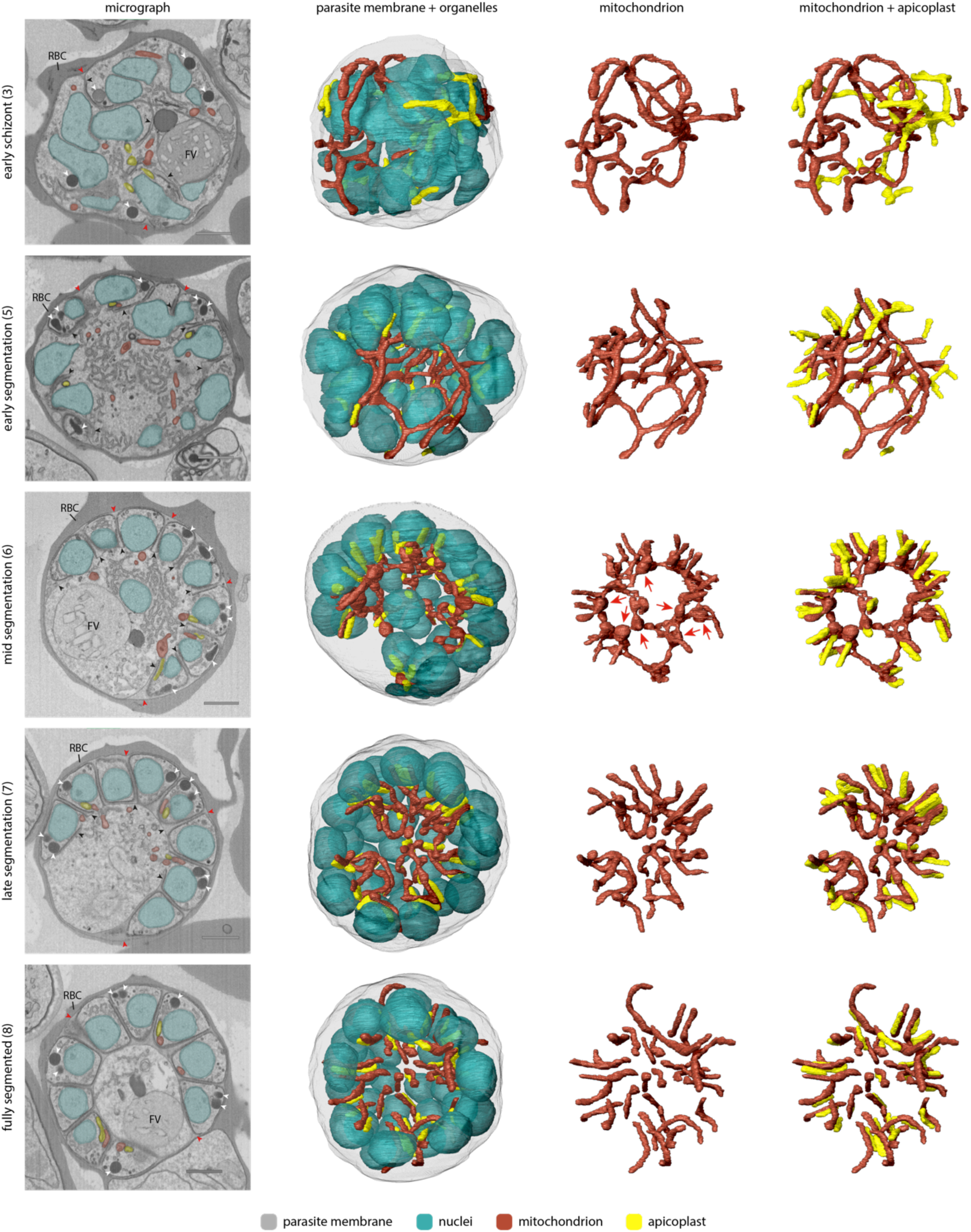
3D rendering of mitochondrion and apicoplast during different stages of schizogony. First column contains representative micrograph images from different schizont stages. The numbers between brackets indicate the parasite ID number and detailed information can be found in table S2 and S3. The red blood cell (RBC) and food vacuole (FV) are indicated by their abbreviations. Rhoptries are indicated by white arrowheads, parasitophorous vacuole membrane is indicated by red arrowheads, and parasite membrane invaginations are indicated by black arrowheads. Scale bars, 1 μm. The second, third, and fourth column contain 3D renderings of parasite membrane (gray, 5% transparency), nuclei (teal, 50% transparency), mitochondrion (red), and apicoplast (yellow). Red arrows indicate merozoite entrance bulins.

**Figure 7.**
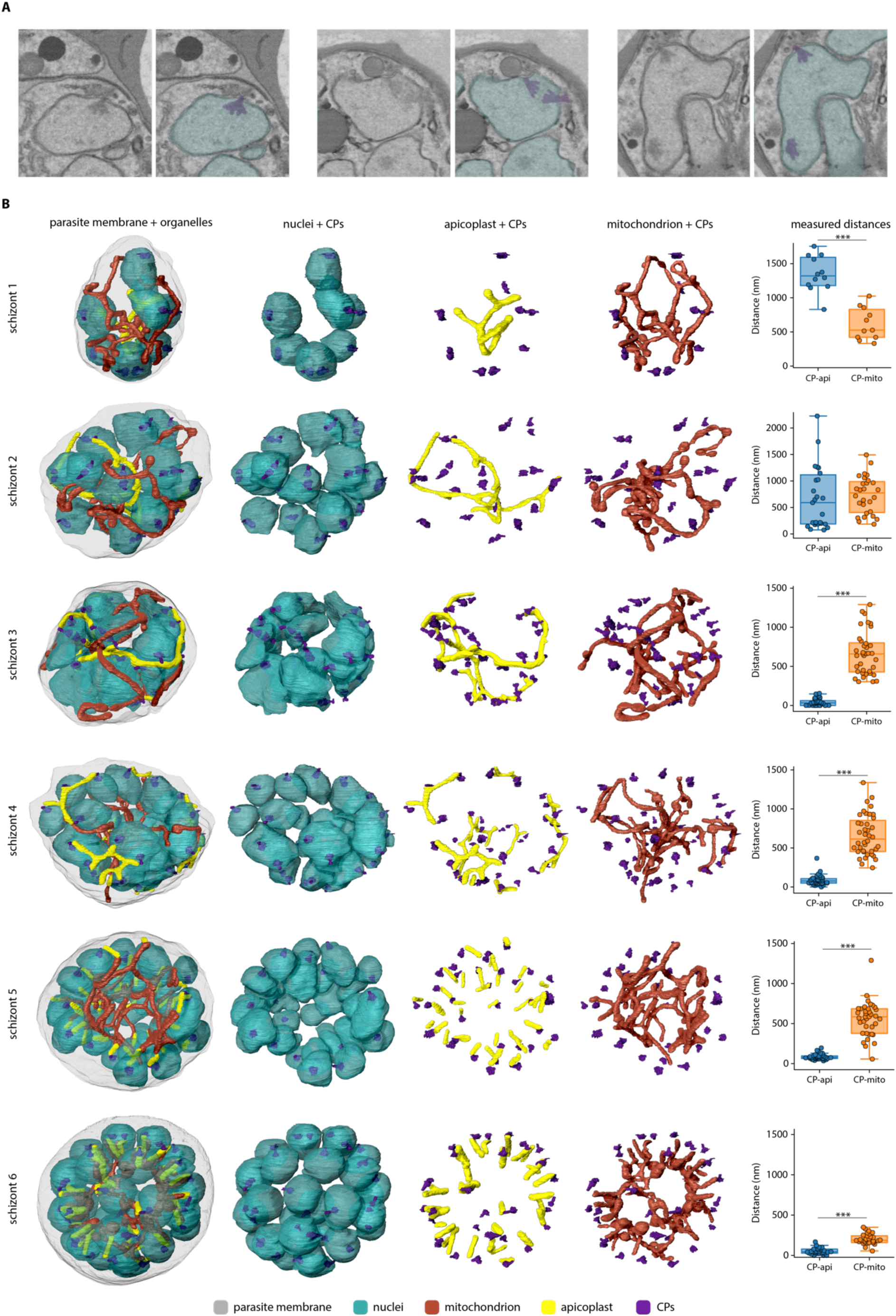
Association of apicoplast and not the mitochondrion with centriolar plaques during schizont development. A) Micrographs of nuclei (teal) with centriolar plaques (CPs, purple). B) 3D rendering of nuclei (teal), apicoplast (yellow), mitochondrion (red), CPs (purple), and parasite membrane (gray, 5% opacity). Parasite ID numbers are indicated on the left side of the micrograph images. Right column shows measured distances between CPs and closest point to the apicoplast or mitochondrion. ***, p < 0.001.

*\* Note: The volumes measured in the FIB-SEM data differ greatly from the volumes measured in the 3D fluorescent microscopy data. This can be explained by the limited spatial resolution of fluorescent microscopy because of the diffraction of light. The diffraction limit of the confocal Airyscan microscope that was used is ∼120 nm in lateral direction and ∼350 nm in axial direction. The diameter of the mitochondrion in asexual blood-stage parasites is ∼140 nm, which is at the edge of the resolution limit. Therefore, the volume measurements of thresholding-based segmentation of the fluorescent signal are not very accurate and quickly over-estimate the volume. These volume estimations should merely be used to compare relative volumes of mitochondrial fragments. The FIB-SEM data has a resolution of 5 nm in lateral direction and 15 nm in axial direction, which allows much more precise visualization of organelles and volume measurements*.

### Interaction between mitochondrion and apicoplast in late stage schizonts

During schizogony, the mitochondrion and apicoplast show different moments of close association. Prior to apicoplast division, the mitochondrion and apicoplast have several apposition sites, which have also been described by Evers *et al.*^20^ (Figure S9A and S9B). It remains unclear if these close associations represent true membrane contact sites that facilitate the exchange of metabolites or lipids between the organelles, or if these are merely random due to the limited space in the parasite. When apicoplast division is finished, the endings of the mitochondrial branches reach towards the basal endings of the apicoplasts (Figure S9D). Subsequently, the branches of the mitochondrial cartwheel structure align with the apicoplast over its entire length (Figure S9C and S9D). This close alignment remains when mitochondrial division is complete.

### Bulbous mitochondrial structures with double membrane invaginations

The parasite mitochondrion does not have a consistent diameter during schizogony. While some parts have a very small diameter other areas of the mitochondrion are more bulbous (Figure S10A). These bulbous parts often contain double membrane invaginations of various size and shape (Figure S10C, S10D, S10F). These bulbous invagination structures (bulins) are found in all schizont stages and vary greatly in shape, size, and location in the mitochondrial network. In early stage schizonts, bulins can be found at branching points and in the middle of a continues branch of the mitochondrial network (Figure S10D, S10F). However, during mid-segmentation stages, when the mitochondrion is oriented in its typical cartwheel structure, bulins are consistently observed at the base of a mitochondrial branch near the merozoite entrance (Figure 6, Figure S10D, S10F). These merozoite-entrance bulins were found in all eight mid-segmentation stage schizonts from two independent experiments (example shown in Movie 9). Bulbous areas at the base of mitochondrial branches are also observed with fluorescent microscopy (Figure S10B). This, and the specific location of bulins, makes them unlikely an artifact of fixation or sample preparation for the FIB-SEM, although this cannot be ruled out completely. Bulins that reside at the entrance of a forming merozoite during the cartwheel phase are typically characterized by contact with the basal end of the divided apicoplast, and a small constriction right above the bulin where the mitochondrial branch enters the merozoite (Movie 10). Bulbous areas at the base of the mitochondrial branches are also observed with fluorescent microscopy (Figure S10B). In late-segmentation schizonts, we observed small bulins at the base of a divided mitochondrial fragment or at the entrance of a merozoite when the mitochondrial branch was not yet divided (Figure S10D, S10F). Sporadically, we also found bulins in the apicoplast (Figure S10E). The significance and function of these bulins remain to be explored, but it is tempting to speculate about a possible role in organelle division.

### Centriolar plaques associate with apicoplast but not mitochondrion during organelle segregation

In mammalian cells, segregation of organelles is coordinated by microtubules that arise from the centrosomes, or so-called microtubule organizational centers (MTOCs). *Plasmodium* parasites lack canonical centrosomes but organize their mitotic spindle from a structure called the centriolar plaque (CP), which is embedded in the nuclear envelope^30,31^. Expansion microscopy studies from Liffner *et al.* have suggested an association of the CPs with both the mitochondrion and the apicoplast during schizogony, suggesting their involvement in organelle segregation^32^. In our FIB-SEM images, we can distinguish the CP by electron dense coffee filter-shaped regions in the nucleus (Figure 7A). In an early schizont that still lacks IMC or rhoptries, nuclei contain one or two CPs, which are oriented to the periphery of the parasite. 3D renderings show no direct association between the CPs and the mitochondrion or apicoplast (Figure 7B, Movie 1). Although the distances between the mitochondrion and CPs (average 616 nm, SD 235 nm) in this early schizont are significantly smaller than the apicoplast-CPs distances (average 1350 nm, SD 260 nm), there is no direct interaction between the mitochondrion and CPs since the smallest CP-mitochondrion distance measured is 332 nm. The significant difference can be explained by the fact that the apicoplast is located in the center of the parasite, while the mitochondrion is larger and stretched throughout the whole cell leading to coincidental closer proximity to the peripheral CPs. In slightly later stage schizonts where IMC and rhoptry formation has started, all nuclei contain either two CPs, or one CP that is dividing. A portion of the CPs associated with the apicoplast, specifically with the endings of apicoplast branches (Movie 2). When the IMC is developed slightly further, all CPs associate with the apicoplast over the total length of the peripherally localized apicoplast network (Movie 3). While two CPs from the same nucleus usually associate with one apicoplast branch (Figure S11A), sometimes these associate with completely different branches of the apicoplast network (Figure S11B). Furthermore, the distances between the CPs and apicoplast are significantly smaller (42 nm average, SD 46 nm) compared to those measured in earlier stages as well as to the unchanged CP-mitochondrion distances (663 nm average, SD 274 nm) (Figure 7B). The association between CPs and apicoplast continues during and after apicoplast division (Movie 4-7). After apicoplast division, each apicoplast fragment is associated with one CP at its peripheral end (53 nm average distance, SD 37 nm) (Movie 5). During mid-segmentation stages, the endings of the mitochondrial branches are close to the CPs (202 nm average distance, SD 65 nm) (Movie 6). However, this seems to be a result of the close association of the mitochondrion with the apicoplast, rather than a direct interaction between the mitochondrion and the CPs. In a fully segmented schizont, the CPs were much smaller and did not show a clear extranuclear compartment (Movie 8). This close and very consistent association between the apicoplast and the CP, suggest an important role in apicoplast segregation, while the mitochondrion likely deploys different mechanisms to accomplish its proper distribution over the forming merozoites.

## Discussion

In contrast to most eukaryotes, the fast-replicating *P. falciparum* asexual blood-stage parasites harbor only a single mitochondrion. Consequently, proper division and distribution of this organelle during schizogony is crucial to ensure all daughter cells receive a mitochondrion. Here, we visualized the poorly understood mitochondrial dynamics in blood and mosquito stages using a new parasite line with a fluorescent mitochondrial marker and super-resolution 3D imaging methods. During blood-stage schizogony, a cartwheel structure is formed and divided into smaller, unequally sized mitochondrial fragments in a stepwise process. Final division into single mitochondria happens during the last stage of merozoite segmentation. These division steps were cross validated by analyzing available FIB-SEM data with nanometer resolution. This also allowed us to reconstruct apicoplast division and its interactions with the mitochondrion. Finally, we showed that the apicoplast but not the mitochondrion associates with the CPs during merozoite formation.

To date, the visualization of *Plasmodium* mitochondria has largely relied on MitoTracker dyes. However, these dyes are toxic at nanomolar concentrations^15,16^ and our data suggest that they may alter mitochondrial morphology (Figure 1). We developed a reporter parasite line harboring a fluorescent mitochondrial marker that shows a more continuous and less punctate staining pattern compared to MitoTracker dyes and is compatible with live and fixed imaging without necessitating antibody labeling. Unfortunately, MitoRed is not well suited for long-term time lapse imaging (>1h) since parasites showed various signs of poor health, probably due to phototoxicity. While expansion microscopy is currently not feasible with MitoRed, addition of a linear epitope tag would make this marker compatible with the required denaturation step.

In line with our earlier observations, we demonstrated multiple mitochondria in gametocyte stages^20^. As discussed by Evers *et al.*, there are several possible reasons for the emergence of multiple mitochondria in gametocytes, such as adaptation to a metabolically varied environment, distribution mechanism of mtDNA, or management of reactive oxygen species. We expand upon these observations by also imaging gametocytes during activation. In males, mitochondria become more dispersed while female mitochondria remain in a tight knot. One possible explanation for this is that mitochondria in males are distributed to specific locations in the cell to provide energy locally for certain processes. In sperm cells, the mitochondrion resides at the base of the flagellum to provide energy for flagellar movement^22^. While we observed close apposition of the mitochondria with axonemal tubulin in some activated males using light microscopy (Fig. S4), this was not consistently observed in all males, and we lack the resolution to prove real association. Another explanation could be that the parasite undergoes a form of mitophagy as a source for proteins, lipids, and nucleotides required for the rapid nuclear division and microgamete formation. Even though mitophagy has not been studied in *Plasmodium*, some homologues of the general autophagy pathway have been identified^33^. Autophagy as a survival mechanism was described for *P. falciparum* and *T. gondii* under starvation conditions^34,35^. In *T. gondii*, the fragmentation of the mitochondrion was reversed by using the established autophagy inhibitor 3-methyl adenine^34^. Alternatively, the distribution of the mitochondria could merely be a consequence of the nuclear expansion in the cell.

Mitochondrial dynamics during mosquito stages is poorly understood and to our knowledge studies have thus far been restricted to *P. berghei*^17,23,24,36^. Here, we visualized the mitochondrion in *P. falciparum* during mosquito stages for the first time. In early oocyst stages, the mitochondrion resembles the extensively branched network from asexual blood-stage schizonts. During oocyst development, the mitochondrial network organizes into multiple MOCs that resemble the cartwheel structure observed in asexual blood stages. Although these mitochondrial observations should be interpreted with care since oocysts did not form salivary gland populating sporozoites and might therefore not be representing healthy oocysts (see the supplement for a more extensive discussion), in *P. berghei* liver-stage schizonts, a very similar mitochondrial organization was observed in sub-compartments created by large membrane invaginations^12,37^. Similar sub-compartments are present during oocyst development^37^. Based on apicoplast visualizations in *P. berghei* and our observations of the formation of MOCs during oocyst stages, mitochondrial and apicoplast dynamics in these sub-compartments in both oocyst and liver stages resemble the dynamics of these organelles in blood-stage schizogony^12,38,39^.

Although the use of new imaging techniques, such as expansion microscopy and 3D volume EM, have revealed new insights in mitochondrial dynamics, many questions about the timing and organization of mitochondrial division remained unanswered^11,32^. In an earlier literature review, we proposed three possible mitochondrial division models: synchronous fission, outside-in fission, or branching point fission^9^. Here, we used a new mitochondrial marker and advanced imaging techniques, such as Airyscan confocal microscopy, to reconstruct mitochondrial fission during schizogony. The use of volume EM provided the resolution required to verify our fluorescence-based mitochondrial fission model while simultaneously shedding light on the division of the second endosymbiotic organelle, the apicoplast. This allows us to propose a new, detailed model of the organellar division and segregation (Figure 8).

**Figure 8.**
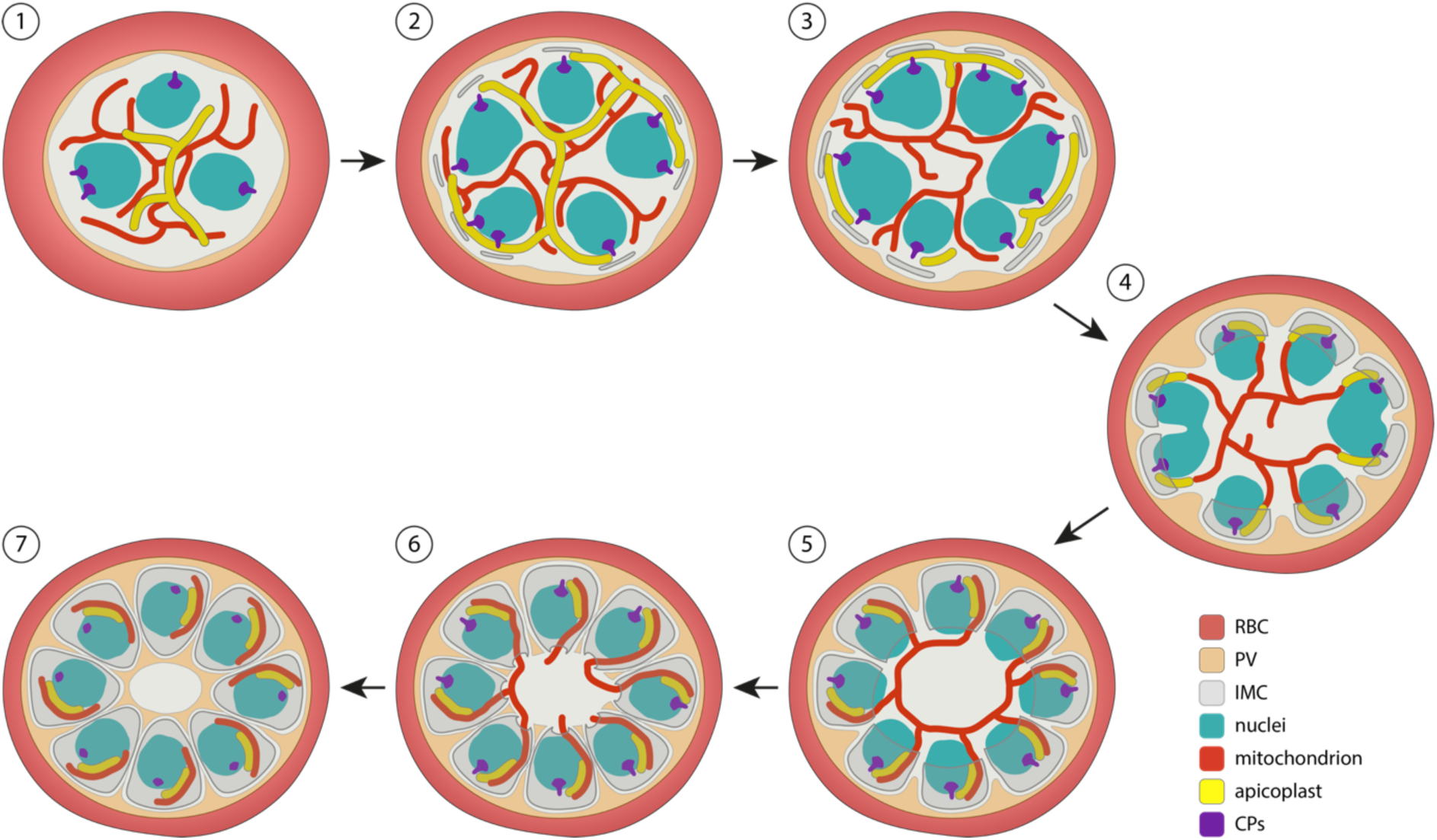
Schematic model for mitochondrial and apicoplast division and segregation in P. falciparum during schizogony. (1) Nuclear division is ongoing, while inner membrane complex (IMC) formation has not started and both the mitochondrion and apicoplast are branched networks. The apicoplast localizes more to the center of the cell, while the mitochondrion is stretched throughout the whole cell. (2) When IMC formation starts, the apicoplast branches associate with the centriolar plaques (CPs) at the periphery of the parasite. (3) The apicoplast divides in a non 2^n^ progression, while it keeps its interaction with the CPs. (4) When nuclear division is finishing, apicoplast division is completely finished. The apical end of the apicoplast fragments associate with the CPs, while mitochondrial branches associate with the basal end of the apicoplast fragments. (5) The IMC develops further and envelops large parts of the nuclei. The mitochondrion orients itself in a cartwheel structure, while its branches align with the apicoplast fragments. (6) IMC formation is almost finished, and just a small opening connects the merozoites to the residual body. The mitochondrion divides in a non 2^n^ progression. The apicoplast still associates with the CPs and aligns with mitochondrial branches/fragments. (7) Merozoite segmentation is complete, the apicoplast loses its clear association with the CPs since they become smaller and do not have a clear extra nuclear compartment anymore. The mitochondrion is fully divided and still aligns with the apicoplast. Red blood cell (RBC), parasitophorous vacuole (PV).

In this model, we describe the cartwheel orientation of the mitochondrion, its non-geometric 2^n^ division, the late timing of division, and its association with the apicoplast. The apicoplast divides when nuclear division is still ongoing and merozoite segmentation has just started. Similar to the mitochondrion, its division does not happen in a geometric 2^n^ progression, but different sized apicoplast fragments are observed in a mid-division stage. The timing and orientation of the apicoplast-mitochondrion appositions, suggest a potential role of the apicoplast in mitochondrial segregation. However, doxycycline treated parasites, that have lost their apicoplast and are chemically rescued by isopentenyl pyrophosphate supplementation, can still produce viable merozoites, suggesting that association with the apicoplast is not essential for mitochondrial segregation^40^. The specific types of membrane contact between both endosymbiotic organelles, whether these consist of direct physical contacts, membrane fusions or tethering proteins, may vary and remain to be explored. However, EM tomography data from Sun *et al*. show there are hints of connecting structures between the mitochondrion and apicoplast in areas where the distance between the organelles is very small and similar to the distance between the inner and outer membranes of the organelles themselves in merozoites, suggesting physical link between the organelles.

In other eukaryotic models, mitochondrial fission is facilitated by adaptor proteins on the cytoplasmic side of the outer mitochondrial membrane that recruit dynamin GTPases, which in turn oligomerize to form a constrictive ring around the organelle^9^. The only conserved adaptor protein in *P. falciparum*, Fission 1 (Fis1), is dispensable in asexual blood stages precluding an essential role in mitochondrial fission during schizogony^41^. Malaria parasites also harbor three dynamin-related proteins^9^. As the timing of the apicoplast division precedes that of the mitochondrion, it is conceivable that (parts of) the same dynamin-based machinery may be reused for mitochondrial fission. Indeed, in a recent pre-print *P. falciparum*, dynamin 2 (*Pf*DYN2) was shown to mediate both apicoplast and mitochondrial fission^42^. Which other proteins comprise the mitochondrial division machinery in *P. falciparum* remains to be explored.

Their morphological features, timing of appearance, and specific location at the entrance of the developing merozoite during mid-segmentation stages suggests that the bulbous invagination structures, or bulins, could play a role in mitochondrial fission. A role in the distribution of certain components – e.g. mitochondrial DNA, proteins, or protein complexes – to the branches of the mitochondrial cartwheel structure that enter the merozoite is also conceivable. However, in earlier stages, bulins are also found at branching points or in continuous parts of the mitochondrion and apicoplast, perhaps suggesting possible roles in more general membrane remodeling of the organellar network.

We also observed CPs, the *P. falciparum* analogue of the centrosome, which function as microtubule organizing centers and are important for mitosis and cell cycle regulation^43,44^. In *T. gondii*, the centrosome associates with the apicoplast and ensures correct segregation of the organelle during daughter cell formation^45,46^. Recent expansion microscopy data suggested interaction of the CP with both the apicoplast and the mitochondrion in *P. falciparum*^32^. From the onset of IMC and rhoptry formation, we observed close apposition of the CPs and apicoplast, but not the mitochondrion (Figure 8). This CP-apicoplast association continues during and after apicoplast division, indicating a role of the CPs in apicoplast segregation. In other apicomplexan parasites, such as *Toxoplasma gondii* and *Sarcocystis neurona*, centrosomes have also been indicated to be involved in apicoplast organization and distribution during cell division^45,47^. The initial absence of CP-apicoplast association and the later association of two CPs from one nucleus with separate apicoplast branches suggests an active recruitment strategy. Motor proteins facilitate intracellular transport of organelles along the cytoskeleton in multicellular eukaryotes. While dynein and kinesin facilitate organelle transport along microtubules, myosin motor proteins transport organelles along actin filaments to specific locations in the cell^48^. Previous studies have shown a critical role of F-actin and myosin F in the inheritance of the apicoplast in *P. falciparum* and *T. gondii*^49–52^. In *T. gondii* parasites that lack myosin F, the apicoplast fails to associate with centrosomes^53^. Therefore, we hypothesize that myosin facilitates recruitment of the apicoplast branches over the actin filaments to the CPs. Although the mitochondrion is close to the CPs in late segmentation stages, the distance is always significantly bigger than the apicoplast-CP distance. Additionally, mitochondrial branches reach much further into the merozoites when fully segmented, compared to the apicoplast. Furthermore, conditional knockout of *PfACT1* (actin-1) did not alter mitochondrial morphology in asexual blood-stage schizogony^49^. Therefore, it remains questionable if the mitochondrial branches are recruited to the CP via a similar mechanism as the apicoplast.

Volume EM is a powerful tool to study biological questions as it allows the visualization of complex, connecting structures and gives spatial and cellular context. Here, we reused available FIB-SEM data, which contains sexual and asexual blood-stage malaria parasites from many different stages of intra-erythrocytic development. Future reinterrogation of the data could facilitate in answering other biological questions that are beyond the scope of this paper, such as rhoptry biogenesis and development of the apical complex.

In this study, we have developed a reporter parasite line harboring a fluorescent mitochondrial marker – integrated in a new genomic locus – that can be used for mitochondrial visualization in blood and early mosquito stages. This allowed us to visualize mitochondrial division in unprecedented detail and describe the relative timing and of mitochondrial fission and segregation using high-resolution confocal microscopy and FIB-SEM image analysis. Combined with new insights in apicoplast division, mitochondrial and apicoplast interactions, and association of the apicoplast with the CP during schizogony, this allowed us to propose a new, detailed model of apicoplast and mitochondrial division during schizogony. These findings pave the way to home in on the molecular mechanisms underpinning mitochondrial and apicoplast division and segregation.

## Materials and Methods

### *P. falciparum* culture and transfections

*P. falciparum* NF54 and MitoRed parasites were cultured in RPMI1640 medium supplemented with 25 mM HEPES, 10% human type A serum and 25 mM NaHCO_3_ (complete medium). Parasites were cultured in 5% human RBCs type O (Sanquin, the Netherlands) at 37°C with 3% O_2_ and 4% CO_2_. For transfection, 60 μg of homology-directed repair plasmid was linearized by overnight digestion, precipitated, and transfected with 60 μg Cas9 plasmid using ring transfection^54,55^. Briefly, a ring-stage sorbitol synchronized parasite culture was transfected with the plasmids by electroporation (310 V, 950 μF). Five hours after transfection, parasites were treated with 2.5 nM WR99210 for five days. Success of transfection was assessed by integration PCR (Fig S1, Table S1).

### Plasmid constructs

To generate the base SIL7 reporter plasmid (pRF0057) the 5’ and 3’ homology regions for SIL7 were amplified from genomic NF54 DNA (Table S1) and cloned into the pBAT backbone^19^ with NgoMIV and AleI (5’) and BmgBI and AatII (3’). For the final MitoRed repair plasmid, first the mScarlet sequence was amplified from p1.2RhopH3-HA-mScarlet (a kind gift from Prof. Alan Cowman)^56^ (Table S1). The mScarlet sequence was cloned into pRF0077 (empty tagging plasmid with PBANKA_142660 bidirectional 3’UTR) with AflII and EcoRI restriction sites, generating pRF0078 intermediate plasmid. The HSP70-3 promotor (prom) and targeting sequence (t.s.) sequence was cloned into pRF0078 with EcoRI and NheI restriction sites, generating pRF0079 intermediate plasmid. The whole mitochondrial marker (HSP70-3 prom + t.s. + mScarlet) was cloned from pRF0079 into pRF0057 with EcoRI and AflII restriction sites, generating pRF0191, the final repair plasmid. CRISPR-Cas9 guide plasmids targeting two different sites in the SIL7 region were generated. Guide oligonucleotides were annealed and cloned into pMLB626 plasmid (a kind gift from Marcus Lee)^57^ using BbsI restriction enzyme, generating the two final guide plasmids (Table S1).

### Growth assay

NF54 WT and MitoRed parasites were synchronized using 63% Percoll centrifugation. Late-stage parasites were isolated from the Percoll gradient and added to fresh RBCs. Four hours later, a 5% sorbitol synchronization was performed, which allowed only young rings that just invaded a new RBC to survive. Ring-stage parasites were counted and three independent cultures of 0.05% parasitemia were set up for each parasite line. Every 24 hours, 10 μl culture was taken and fixed in 100 μl 0.25% glutaraldehyde in PBS up until day 5. To prevent overgrowth, parasite cultures were cut back 1/50 after samples were taken on day 3. Before readout, fixative was taken of, and parasite DNA was stained with 1:10,000 SYBR Green in PBS. Parasitemia was determined by measuring SYBR Green positive cells with a Cytoflex flow cytometer (Beckman Coulter Cytoflex) using the 488 nm laser. Final parasitemia on day 4 and 5 was adjusted for the 1/50 dilution factor, explaining why final parasitemia can reach more than 100%.

### Immunofluorescence assays and fixed fluorescence imaging

Immunofluorescence assays were performed on asexual and sexual blood-stage parasites, using the same fixation and staining protocols. Asexual blood-stage parasites were usually synchronized with 5% sorbitol to get them in the preferred stage for imaging. For tight synchronization, late-stage parasites were isolated with 63% Percoll centrifugation and added to fresh RBCs. Four hours later, a 5% sorbitol was performed to select for young rings. Parasites were settled on a poly-L-lysine coated coverslip for 20 min at 37°C. Parasites were fixed (4% EM-grade paraformaldehyde, 0.0075% EM-grade glutaraldehyde in PBS)^58^ for 20 min and permeabilized with 0.1% Triton X-100 for 10 min. Samples were blocked with 3% bovine serum albumin (BSA) (Sigma-Aldrich) in PBS for 1 h. Primary and secondary antibody incubations were performed for 1 h in 3% BSA. The nucleus was visualized by staining with 1 μM DAPI in PBS for 1 h. PBS washes were performed between different steps. Parasites were mounted with Vectashield (Vector Laboratories). Images were taken with a Zeiss LSM880 or LSM900 Airyscan microscope with 63x oil objective with 405, 488, 561, 633 nm excitations. Images were Airyscan processed before analysis with FIJI software. MitoTracker stainings (including MitoTracker™ Orange CMTMRos, Red CMXRos, Deep Red FM, all from Thermo Fisher Scientific) were done before settling and fixation by incubation of the parasites with 100 nM MitoTracker for 30 min at 37°C, followed by a wash with complete media. The IMC protein GAP45 was labeled using the anti-GAP45 rabbit antibody (1:5000) (a kind gift from Julian Rayner)^27^ and goat anti-rabbit AlexaFluor 488 antibody (1:500, Invitrogen). Alfa-tubulin was labeled with an anti-alfa tubulin mouse antibody (1:500, Thermo Fisher Scientific) and chicken anti-mouse AlexaFluor 488 antibody (1:400, Invitrogen). 3D visualization and quantifications were done in Arivis 4D Vision software. For mitochondrial measurements, threshold-based segmentation was used. For nuclei, blob-finder function was used for segmentation. Number of segmented objects and volume of objects was determined automatically in Arivis software.

### Gametocyte generation and mosquito feeds

Gametocyte cultures were maintained in a semi-automatic culturing system with media changes twice a day^59^. MitoRed gametocytes used for imaging were induced by Albumax supplementation. A mixed asexual blood-stage culture of 1% was set up and maintained in medium supplemented with 2.5% Albumax II (Thermo Fisher Scientific) without human serum for four days^60^. After four days, parasites were cultured in complete medium again for further gametocyte development. For mosquito feeding, MitoRed gametocytes were stress induced through asexual overgrowing. A mixed asexual blood-stage culture of 1% was set up and maintained for 2 weeks. At day 15 after gametocyte induction, gametocytes were fed to *Anopheles stephensi* mosquitoes (Sind-Kasur Nijmegen strain)^61^. 24 hours after feeding, several mosquitoes were dissected, and blood bolus was obtained for live imaging of ookinetes.

### Live imaging of asexual blood-stage parasites

Sorbitol-synchronized MitoRed and NF54 schizonts were stained with 0.5 μg/ml Hoechst 33342 (Invitrogen, H3570) for 30 min at 37°C for nuclear staining. MitoTracker stainings (including MitoTracker™ Orange CMTMRos, Red CMXRos, Deep Red FM, Rhodamin123, all from Thermo Fisher Scientific) were done by incubation of the parasites with 100 nM MitoTracker or 1 μg/ml Rhodamin123 for 30 min at 37°C, followed by a wash with complete medium. Stained parasites were diluted 1:40 in complete medium and settled for 20 min at 37°C in a poly-L-lysine coated μ-slide 8-well imaging chamber (Ibidi). Unbound cells were washed away with phenol red free complete medium in which cells were also kept during imaging. Parasites were imaged on a Zeiss LSM880 Airyscan microscope with 37°C heated stage table and 63x oil objective. Images were Airyscan processed before analysis with FIJI software.

### Live imaging of mosquito-stage parasites

Ookinetes were obtained from the blood bolus of infected mosquitoes 24 hours after feeding. Ookinetes were stained by mouse monoclonal anti-*Pf*s25 conjugated antibody (made in house, final concentration 15µg/ml). Stained sample was applied on a glass slide and covered with a glass coverslip. The sample was immediately imaged on a Zeiss LSM880 Airyscan microscope with 63x oil objective. Mosquito midguts were dissected at day 7, 10, and 13 after infection and put on a glass slide in PBS covered with a glass coverslip. Samples were imaged immediately on a Zeiss LSM880 or LSM900 microscope with 63x oil objective. Oocysts were identified based on their fluorescent mitochondrion and round shape in the brightfield channel. All images were Airyscan processed before analysis with FIJI software. 3D segmentations and visualizations were done by manual thresholding of the fluorescent signal in Arivis 4D vision software. Salivary glands were dissected on day 13, 16, and 21 after infection and stained with mouse monoclonal anti-CSP conjugated antibody (made in house, final concentration 1µg/ml). Stained glands were applied on a glass slide and covert with a glass coverslip. Samples were imaged on a Zeiss Axioscope A1 microscope with AxioCam ICc1.

### FIB-SEM image analysis

FIB-SEM image stacks were reused from Evers *et al.*^20^ (EMPIAR-10392). For these stacks, gametocyte-induced iGP2 parasite cultures were MACS purified. During this process many late-stage asexual parasites in these cultures were co-purified and fixed in the agarose blocks used for FIB-SEM imaging. Detailed sample preparation and FIB-SEM imaging methods are described in Evers *et al.*^20^. All image processing, visualizations and analysis was done in ORS Dragonfly software (2022.2). Segmentations were done by either manual segmentation or deep learning-based segmentation. Deep learning-based segmentations were manually reviewed and corrected when necessary. 3D renderings of segmented regions were converted to triangle meshes for visualization.

## Acknowledgements

We thank members from the molecular malaria research group for the discussions. We also thank Aniek Garritsen for her contributions to the generation of the SIL plasmids. We do greatly appreciate the help from Chiara Andolina and Nicholas Proellochs with mosquito experiments. We would like to thank Astrid Pouwelsen, Jolanda Klaassen, Laura Pelser-Posthumus, Saskia Mulder and Jacqueline Kuhnen for breeding of mosquitoes and handling of the infected mosquitoes. We thank the Radboud Technology Center Microscopy, Radboud Technology Center flowcytometry, and Radboudumc Electron Microscopy Center for the use of their facilities. We are grateful to Alan Cowman for sharing the p1.2RhopH3-HA-mScarlet plasmid and Julian Rayner for sharing the anti-GAP45 antibody. J.M.J.V. is supported by an individual Radboudumc Master-PhD grant. A.B.V. is supported by an NIH grant (R01 AI028398).

