## Supplementary Information 1 for "Detailing organelle division and segregation in *Plasmodium falciparum*"

**Supplemental information S1.**

**Selection of new genomic integration sites**
A previous study has described successful integration of a reporter gene into a so-called silent intergenic locus (SIL) in *P. berghei*[1]. We extended the same concept to *P. falciparum* and identified potentially suitable chromosomal breakpoints in *P. falciparum* conserved between *Plasmodium vivax* and the rodent malaria parasite lineages[2]. By using these SIL sites, we prevent the need for additional gene replacements commonly used to introduce transgenes. For instance, NF54HT-GFP-luc parasites had the GFP-luc cassette introduced in the presumed silent *Pf*s47 locus[3]. Meanwhile, it has been demonstrated that this gene can play an important role in mosquito infections[4,5]. Another commonly used integration site is the *PfRH3* pseudogene, for instance for integration of a dimerizable Cre gene[6,7]. Although *PfRH3* is transcribed and not translated in blood stages, the RH3 protein has been detected in sporozoites[8,9]. The genes flanking the SIL sites have been rearranged during the evolution of *P. falciparum* and we reasoned that there would not be any genetic constraints to keep these genes physically linked. We analyzed available data on gene expression and potential transcription start sites to target a locus that lacks any detectable (regulation of) gene expression in the *Plasmodium* life-cycle stages[10]. In addition, UTRs of the flanking genes need to remain intact.

Applying the combined criteria, we identified three loci on *P. falciparum* chromosomes 7, 12, and 14. The site on chromosome 12 (genomic location between PF3D7_121220 and PF3D7_1212100) proved very difficult to clone, with frequent plasmid rearrangements and very low plasmid yields. A total of 3 transfections targeting the site on chromosome 14 (genomic location between PF3D7_1438100 and PF3D7_143800) did not result in any transgenic parasite lines, suggesting either technical challenges or some important function of this locus in ABS parasite viability. The third SIL, termed SIL7 as it is on chromosome 7, locates between PF3D7_0715900 and PF3D7_0716000 and was used for integration of the mitochondrial marker. We generated an integration plasmid containing 5’ and 3’ homology regions (HRs) and the mitochondrial marker (Figure S1A).

**Considerations for the use of SIL7 for stable transgene integration**
Integration of the fluorescent mitochondrial marker in SIL7 did not affect parasite growth or development in blood and mosquito stages up until oocyst formation. Although we did observe several free sporozoites in our dissected mosquito samples, we never observed sporozoites in the salivary glands. This suggests that sporozoites might have a developmental defect that prevents them from populating the salivary glands. One possible explanation could be the presence of the fluorescent mitochondrial marker, which might be toxic for this stage specifically. Even though we used an organelle-specific promoter, the high, HSP70-3 (PF3D7_1134000) promoter driven mito-mScarlet expression combined with limited resources, and possible lack of feedback loops to control protein levels could be overwhelming the small sized mitochondrion. Alternatively, integration in SIL7 might be disruptive for sporozoite development, possibly due to interference with genetic or epigenetic processes in this stage. Therefore, we conclude that SIL7 is an excellent integration site when studying blood or mosquito stages up until oocyst development, but it might not be well suited to study sporozoite or liver stages.

**Table S1. Primer and guide RNA sequences for generation of repair and guide plasmids.**

| Primer name | Primer function | Sequence | Restriction site |
|---|---|---|---|
| Generation repair plasmids | | | |
| JV069 | HSP70-3 prom + t.s. F | AATAAAGAATTCTTGCATGCCCCATAATTTTCAC | EcoRI |
| JV070 | HSP70-3 prom + t.s. R | ATTAAAGCTAGCAGCATCTTCATCATATTTTCTACC | NheI |
| JV071 | mScarlet F | AATAATGAATTCAAAGCTAGCATGGTGAGCAAGGGCGAGG | EcoRI, NheI |
| JV072 | mScarlet R | AATAATCTTAAGTTACTTGTACAGCTCGTCCATGC | AflII |
| NF001 | SIL7 5' HR F | AAAGCCGGCGTCGACGAAAAAAAAGAAGAGTAGAGCAGTAC | NgoMIV |
| NF002 | SIL7 5' HR R | TAACACGTACGTGAGGTAATATAACATTGAATTTATAATACAT TAC | AleI |
| NF003 | SIL7 3' HR F | AAACCACGTCTTGGAAATGTGTAGCACTTTTTTCATTCC | BmgBI |
| NF004 | SIL7 3' HR R | AAAGACGTCGTCGACCCTATAAAATAAAAATGATTCCAACAAA AAAG | AatII |
| Guide RNA sequences | | | |
| Pf0084 | SIL7 guide 1 sense | TATTGTATATGTGGTAATAAATAAA | |
| Pf0085 | SIL7 guide 1 antisense | AAACTTTATTTATTACCACATATAC | |
| Pf0086 | SIL7 guide 2 sense | TATTGATTCAATATAATAAGGTCAA | |
| Pf0087 | SIL7 guide 2 antisense | AAACTTGACCTTATTATATTGAATC | |
| Integration PCR | | | |
| NP297 | Integration PCR 5' F | GCTCACCTTAAATGTTCCAC | |
| JV104 | Integration PCR 5' R | ATTATATGTGAAAATTATGGGGCATGC | |
| NP190 | Integration PCR 3' R | AGTCATATCCAGGAATAAACATAC | |
| NP298 | Integration PRC 3' R | CGTTCATGCTTTCACAAGAAC | |

***Table S1. Primer and guide sequences for generation of repair and guide plasmids.*** *Used abbreviations: HR = homology region, F = forward primer, R = reverse primer. Overhang for restriction sites are red, restriction sites are underlined, and gRNA sequences are blue.*

| Parasite ID | Nr of nuclei | Total nuclei volume (μm³) | Average volume nuclei (±SD) (μm³) | Nr of mito fragments | Total mito volume (μm³) | Average volume mito fragment (±SD) (μm³) | Nr of apicoplas t fragments | Total apicoplast volume (μm³) | Average volume apicoplast fragment (±SD) (μm³) | Nr of CPs | Total parasite volume (μm³) |
|---|---|---|---|---|---|---|---|---|---|---|---|
| Schizont 1* | 8 | 14.05 | 1.76 (0.31) | 1 | 1.29 | 1.294 | 1 | 0.57 | 0.573 | 12 | 60.11 |
| Schizont 2 | 15 | 18.26 | 1.22 (0.27) | 1 | 1.06 | 1.055 | 1 | 0.37 | 0.372 | 29 | 74.29 |
| Schizont 3 | 20 | 24.42 | 1.22 (0.21) | 1 | 1.44 | 1.443 | 1 | 0.71 | 0.711 | 36 | 84.28 |
| Schizont 4 | 23 | 27.48 | 1.19 (0.33) | 1 | 1.33 | 1.334 | 8 | 0.51 | 0.064 (0.043) | 40 | 98.63 |
| Schizont 5 | 32 | 22.19 | 0.69 (0.24) | 1 | 1.09 | 1.093 | 36 | 0.31 | 0.009 (0.002) | 36 | 81.92 |
| Schizont 6 | 32 | 19.22 | 0.60 (0.04) | 1 | 1.19 | 1.193 | 31 | 0.52 | 0.017 (0.005) | 32 | 86.14 |
| Schizont 7 | 34 | 17.20 | 0.51 (0.02) | 21 | 0.84 | 0.040 (0.026) | 35 | 0.42 | 0.012 (0.002) | 34 | 72.41 |
| Schizont 8 | 32 | 18.11 | 0.57 (0.02) | 32 | 0.86 | 0.027 (0.005) | 32 | 0.35 | 0.011 (0.002) | 32 | 78.63 |

***Table S2. Description of segmented schizonts used in this paper.***
*Parasite is not complete so displayed numbers might not represent the total numbers/volumes in this cell.

| Parasite ID | Detailed description | | Representative micrograph |
|---|---|---|---|
| Schizont 1 | **Nuclei** | Early stage schizont with 8 nuclei, nuclear division is still ongoing, nuclei are large and irregularly shaped. | 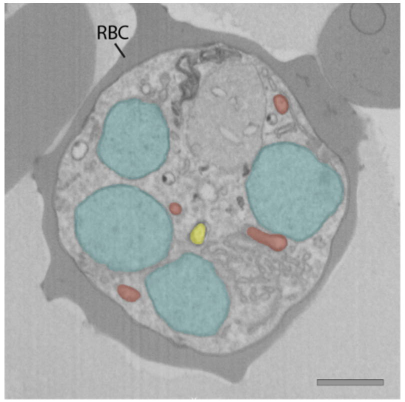  |
| | **IMC and rhoptries** | No IMC formation and no rhoptries. | |
| | **Apicoplast** | The apicoplast is smaller and less complex than the mitochondrion and locates to the center of the parasite. | |
| | **Mitochondrion** | The mitochondrion is one large network stretched throughout the whole cell. | |
| | **CPs** | CPs are dividing and do not interact with the apicoplast or mitochondrion. | |
| Schizont 2 | **Nuclei** | Schizont has 15 nuclei, nuclear division is still ongoing, nuclei are large and irregularly shaped. | 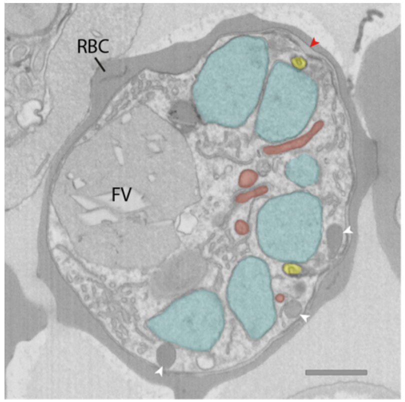  |
| | **IMC and rhoptries** | IMC and rhoptry formation has started (one larger, dark rhoptry and one smaller, light rhoptry per pair). | |
| | **Apicoplast** | Apicoplast branches are more elongated, and some branches associate with a portion of the CPs. | |
| | **Mitochondrion** | The mitochondrion is one large network stretched out throughout the whole cell. | |
| | **CPs** | CPs are dividing or have divided and are localizing further apart from each other within one nucleus. Some CPs interact with the apicoplast branches. | |

***Table S3. Detailed textual description of segmented parasites used in this paper.*** *RBC and food vacuole (FV) are indicated by abbreviations. Rhoptries (white arrowhead), parasitophorous vacuole membrane (red arrowhead) and parasite membrane invagination (black arrowhead), nuclei (teal), mitochondrion (red), apicoplast (yellow).*

| Parasite ID | Detailed description | | Representative micrograph |
| --- | --- | --- | --- |
| Schizont 3 | **Nuclei** | Schizont has 20 nuclei, nuclear division is still ongoing, nuclei are large and irregularly shaped. | 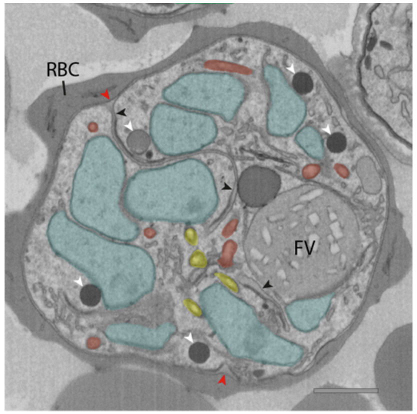  |
| | **IMC and rhoptries** | Large parasite membrane invaginations, below which IMC is being formed, IMC does not yet show curvature, rhoptries are present (one larger, dark rhoptry and one smaller, light rhoptry per pair). | |
| | **Apicoplast** | Apicoplast has developed more smaller branches compared to earlier stages, the apicoplast associates with all CPs. Most of the apicoplast branches located to the periphery of the parasite. | |
| | **Mitochondrion** | The mitochondrion is one large network stretched out throughout the whole cell. | |
| | **CPs** | CP division is finished and CPs are located further apart from each other within one nucleus. | |
| Schizont 4 | **Nuclei** | Schizont has 23 nuclei, nuclear division is still ongoing, most nuclei are large and irregularly shaped and are still dividing, some are smaller and nuclear division seems to be finished (contain only one CP). | 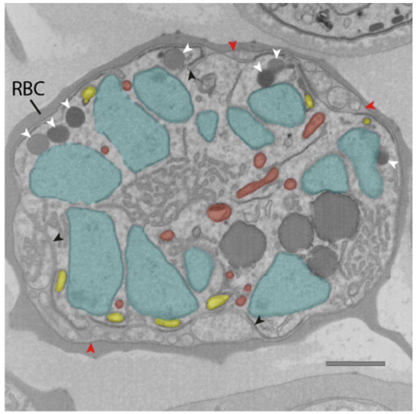  |
| | **IMC and rhoptries** | Parasite membrane starts to curve around the forming merozoites, IMC is showing clear curvature, rhoptries are present below the IMC (the rhoptries from a pair are similar size, but one rhoptry is darker and the rhoptry neck is elongated further than the lighter rhoptry). | |
| | **Apicoplast** | Apicoplast is mid division and consists of 8 different-sized fragments. The apicoplast localizes completely to the periphery of the parasite. The apicoplast interacts with all CPs and each ending of an apicoplast branch is interacting with a CP. | |
| | **Mitochondrion** | The mitochondrion is one large network stretched out throughout the whole cell. | |
| | **CPs** | CP division is finished and larger nuclei that are still undergoing division contain two dispersed CPs, while the smaller nuclei that seem to have finished division contain one CP that localizes close to the apical end of the forming merozoites. | |

| Parasite ID | Detailed description | | Representative micrograph |
|---|---|---|---|
| Schizont 5 | **Nuclei** | Schizont has 32 nuclei, nuclear division is almost finished and only four nuclei are still undergoing nuclear division (they contain two dispersed MTOCs and are larger). | 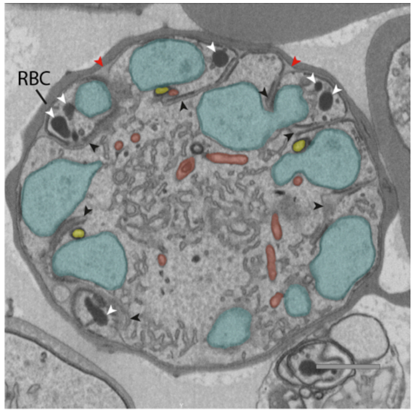 |
| | **IMC and rhoptries** | Parasite membrane is curved further around the forming merozoites, IMC has developed further and is enveloping part of the nucleus. Rhoptries are present below the IMC (rhoptries of one pair now both have the same dark color and rhoptry neck has elongated, although one rhoptry is still longer than the other). | |
| | **Apicoplast** | Apicoplast division has finished and each apical apicoplast ending is associating with a CP. | |
| | **Mitochondrion** | The mitochondrion starts to orient itself in a cartwheel structure where the branches are entering the forming merozoites and are often interacting with the basal end of the apicoplast fragments. | |
| | **CPs** | CP division is finished, and most nuclei contain only one CP that is localized to the apical end of the forming merozoite. | |
| Schizont 6 | **Nuclei** | Schizont has 32 nuclei and nuclear division is completely finished, all nuclei are small, round shaped and contain one CP. | 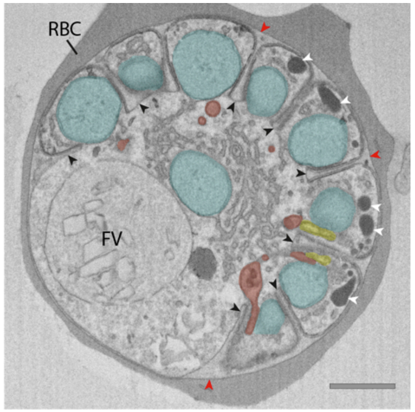 |
| | **IMC and rhoptries** | IMC has developed further and envelops a large part the nucleus. Rhoptries are present at the apical end and seem to be fully developed (rhoptries from one pair have the same shape and dark color and have an elongated rhoptry neck). | |
| | **Apicoplast** | The apicoplast is completely divided and each apical apicoplast ending associates with a CP. | |
| | **Mitochondrion** | The mitochondrion is oriented in a clear cartwheel structure where the branches are entering the forming merozoites and align with the apicoplast. | |
| | **CPs** | CP division is finished, and each nucleus contains one CP that is located to the apical end of the forming merozoite. | |

| Parasite ID | Detailed description | | Representative micrograph |
|---|---|---|---|
| Schizont 7 | **Nuclei** | Late schizont with 34 nuclei. Nuclear division is completely finished, all nuclei are small, round shaped and contain one CP. | 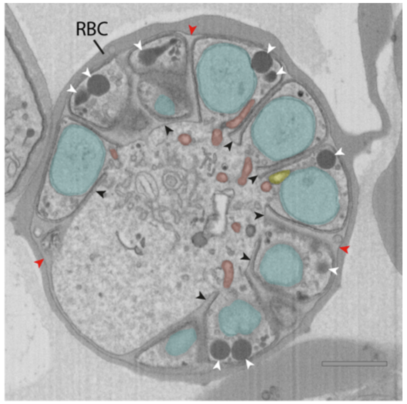 |
| | **IMC and rhoptries** | IMC formation seems almost completely finished and IMC envelops the complete nucleus. Merozoites have a small basal opening through which the mitochondrion enters the parasite. | |
| | **Apicoplast** | The apicoplast is completely divided and each apicoplast ending is associate with a CP. | |
| | **Mitochondrion** | The mitochondrion is dividing and consists of 21 fragments of different sizes. The mitochondrial branches and fragments that enter the merozoite completely align with the apicoplast. | |
| | **CPs** | CP division is finished, and each nucleus contains one CP that is located to the apical end of the forming merozoite. | |
| Schizont 8 | **Nuclei** | Late stage schizont with 32 nuclei. Nuclear division is finished, all nuclei are small, round shaped and contain one CP. | 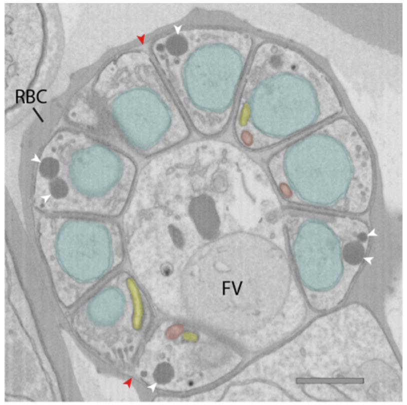 |
| | **IMC and rhoptries** | IMC formation is almost finished and most merozoites are completely enveloped by parasite membrane and pinched of from the residual body. Some merozoites are still connected to the basal body and have a small opening at their apical end. | |
| | **Apicoplast** | The apicoplast is completely divided and does not clearly associate with the CP anymore. | |
| | **Mitochondrion** | The mitochondrion has completely divided and consists of 32 fragments. Each mitochondrial fragments localizes fully into a merozoite, except for two mitochondrial fragments that still enter the residual body through the small apical opening. Mitochondrial fragments align with the apicoplast fragments and are slightly longer. | |
| | **CPs** | CP division is finished, and each nucleus contains one CP that is located to the apical end of the forming merozoite. CPs don't have an extra nuclear compartment and are smaller compared to earlier stages. | |

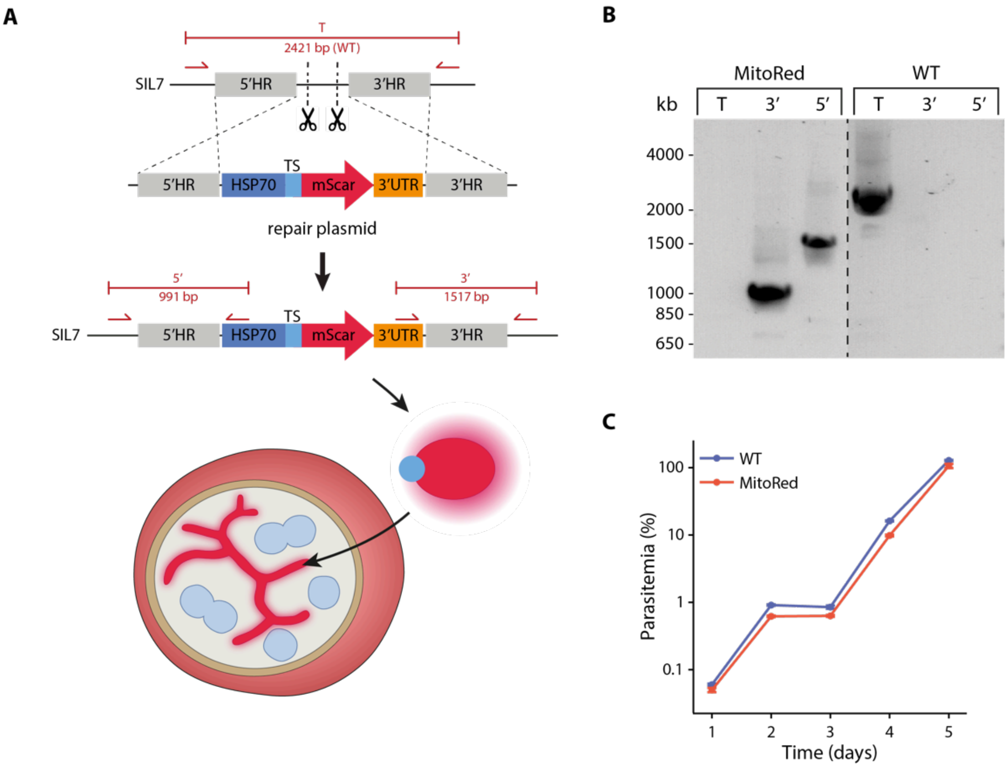


***Figure S1. Generation and verification of MitoRed parasite line.*** *A) Schematic overview of strategy to generate a parasite line harboring a fluorescent mitochondrial marker. CRISPR-Cas9 is used to create two double-strand breaks at SIL7 (indicated by scissors). A construct containing the promotor and targeting sequence of the mitochondrial protein HSP70-3 (PF3D7_1134000) fused with mScarlet is integrated by double homologous recombination. Once integrated, the mitochondrial targeted mScarlet is expressed and led to fluorescent staining of the mitochondrion. B) Diagnostic PCR of MitoRed parasite line with WT- and integration-specific primer combinations (indicated in panel A) demonstrating successful 5' and 3' integration and the absence of WT parasites in the MitoRed line. C) Growth assay showing similar growth of MitoRed and WT parasites. Three independent cultures were set up from one tightly synchronized parasite culture for both MitoRed and WT. Samples were taken over a 5-day period and parasitemia (corrected for dilution factors) was determined with flow cytometry. Error bars (note they are quite small) indicate standard deviation.*

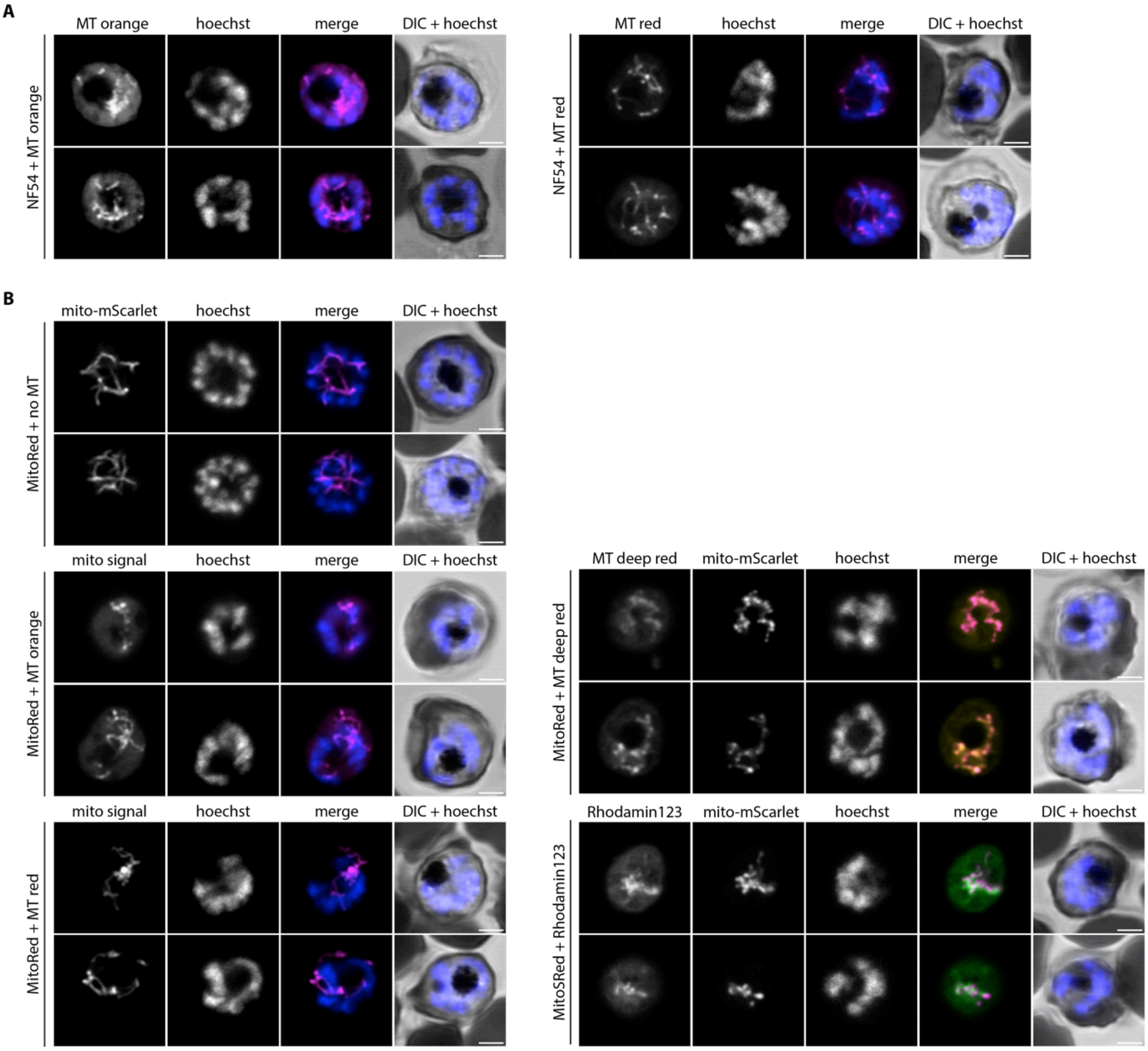


***Figure S2. Comparison of MitoTracker and the mitochondrial marker for live fluorescence imaging.*** *A) live imaging of WT parasites stained with MitoTracker Orange CMTMRos (MT orange) or MitoTracker Red CMXRos (MT red). B) Live imaging of MitoRed stained with MT orange, MT red, MitoTracker Deep Red FM (MT deep red), Rhodamin123 or without staining. Mito signal is the combined MitoTracker and mito-mScarlet signal that is observed in this channel. Parasites were stained with Hoechst 33342 to visualize DNA. All images are single slices of Z-stacks taken with Airyscan confocal microscope. Scale bars, 2 μm.*

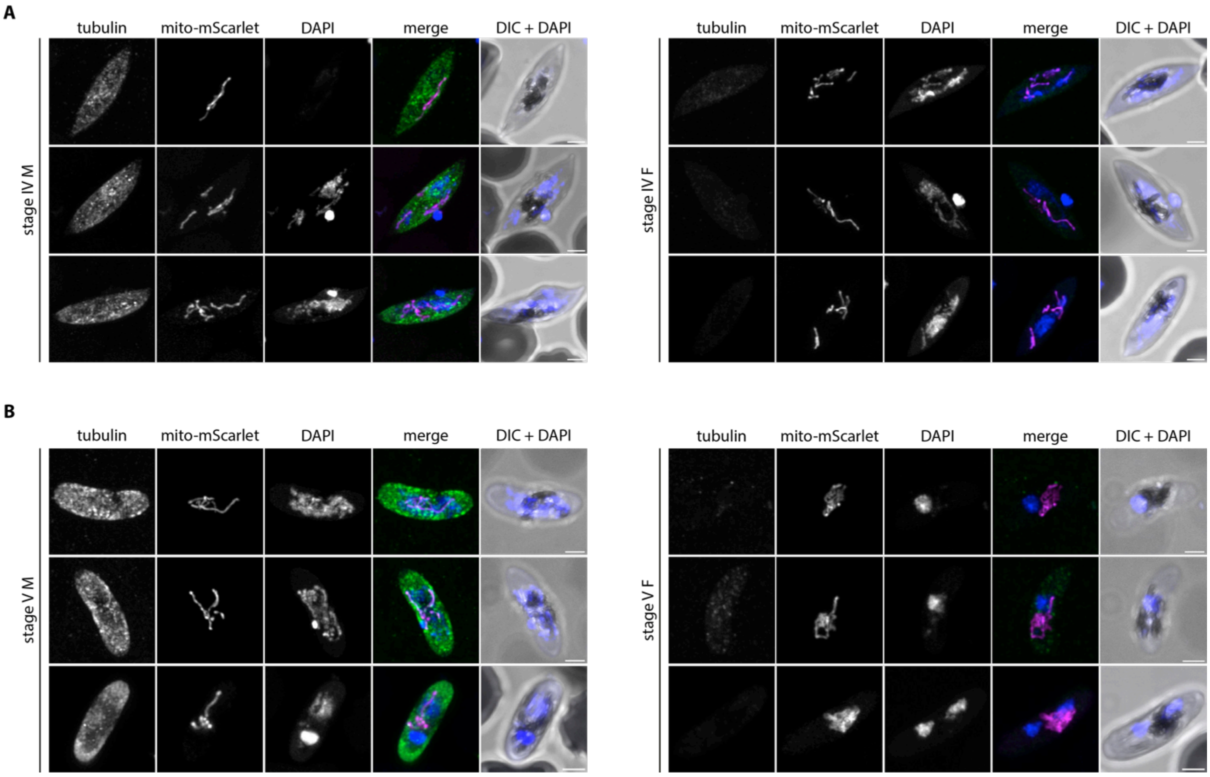


***Figure S3. Mitochondrial morphology in stage IV and V male and female gametocytes.*** *Immunofluorescence assay on male and female MitoRed gametocytes stage IV (A) and stage V (B), stained with anti-β-tubulin (green) and DAPI (blue). Male (M) and female (F) gametocytes are distinguished based on the intensity of the tubulin signal (males high, females low). Images are maximum intensity projections of Z-stacks taken with an Airyscan confocal microscope. Scale bars, 2 μm.*

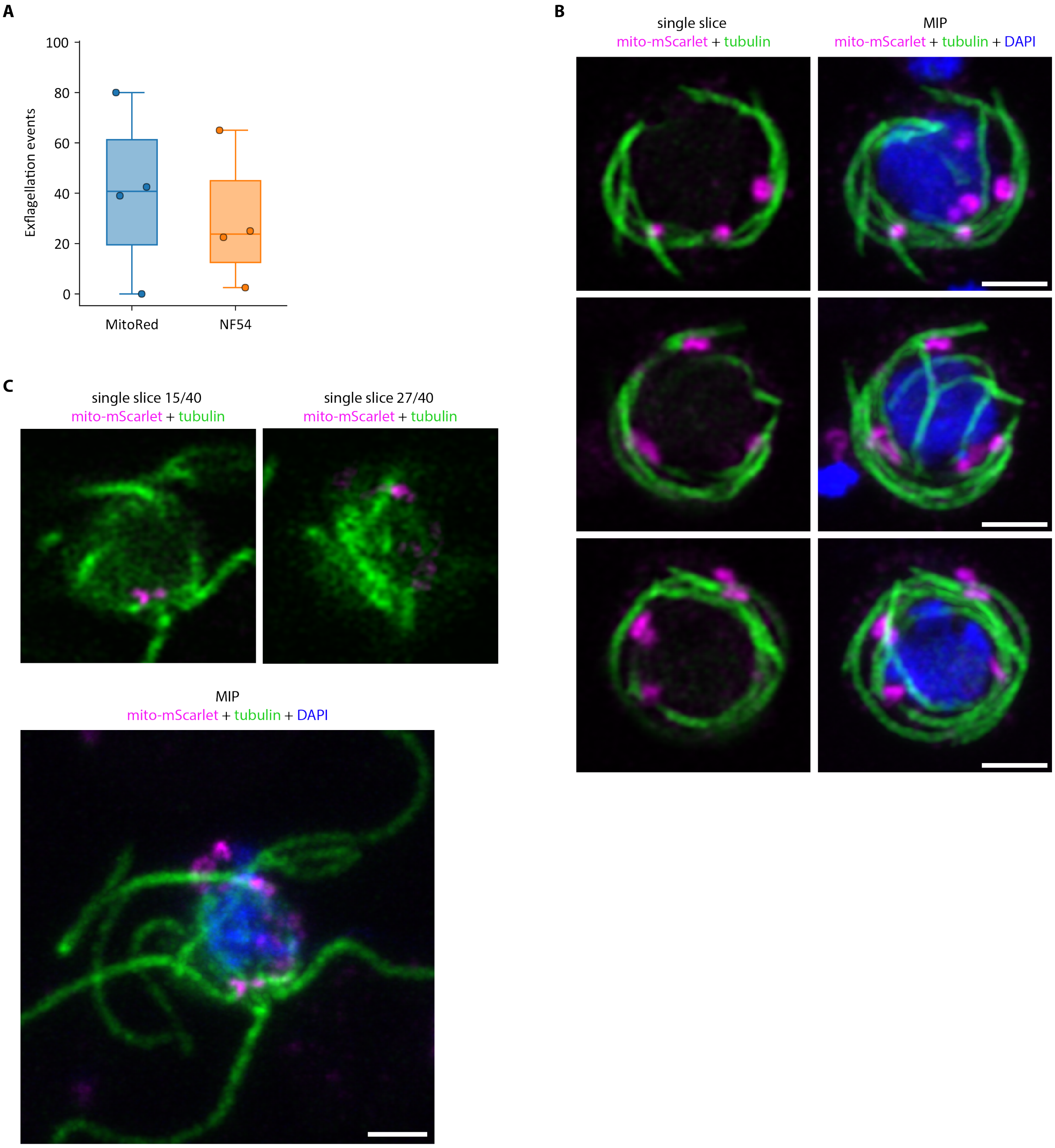


***Figure S4. Mitochondrial association with axonemes in activated male gametocytes.*** *A) Exflagellation events in MitoRed and NF54 parasites 20 min after activation in four cultures in two independent experiments. Unpaired t-test showed no significant difference. B) Three examples of activated males where the mitochondria (magenta) localize closely to the axonemal tubulin (green) in MitoRed parasites. Left images are single slices, right images are maximum intensity projections (MIPs). C) Exflagellating MitoRed male gametocyte with apposition of the mitochondria with the axonemal tubulin. Top images are single slices and crops from bottom image, which is a MIP. Scale bars, 2 μm.*

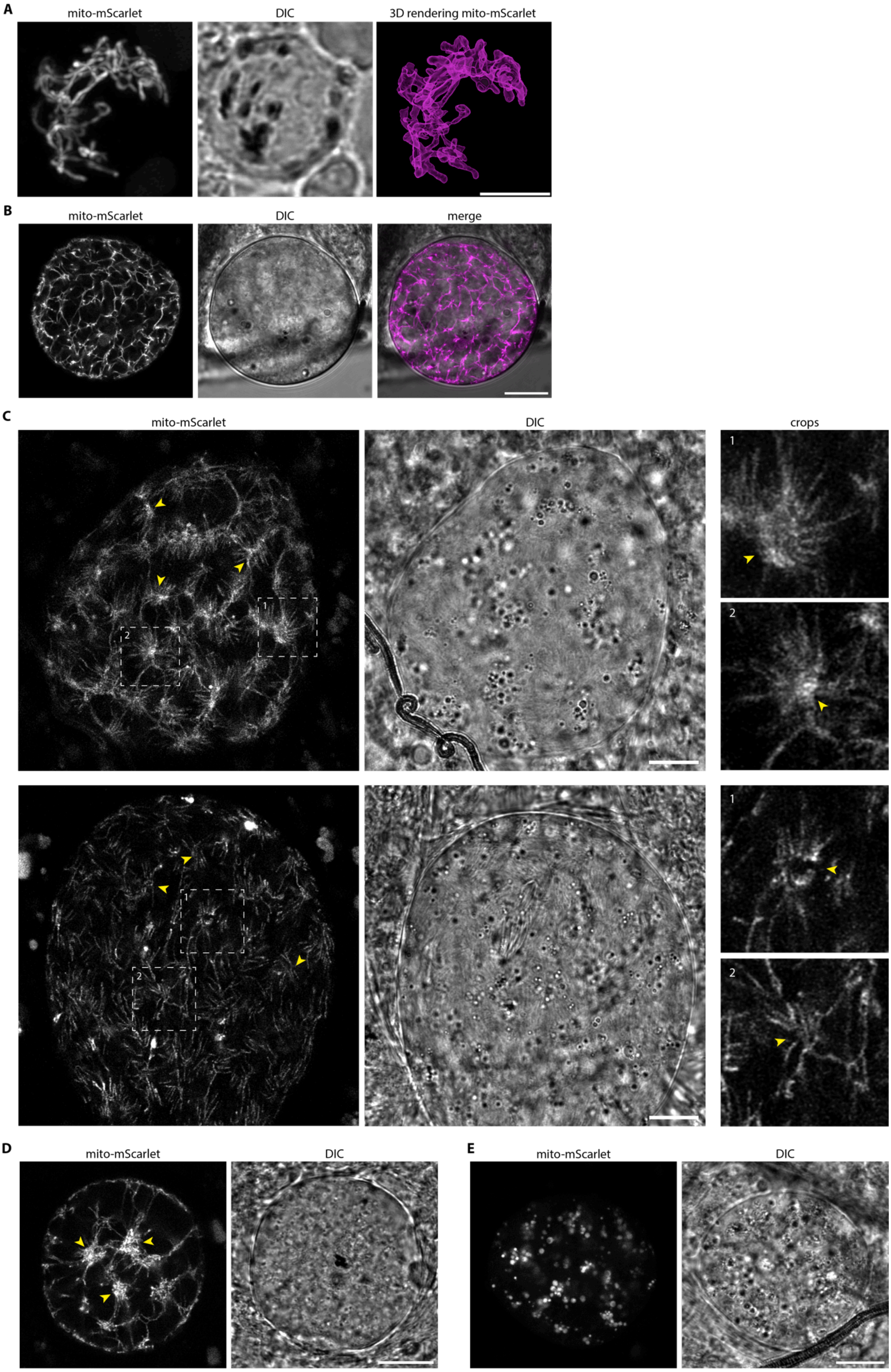
A
mito-mScarlet
DIC
3D rendering mito-mScarlet
B
mito-mScarlet
DIC
merge
C
mito-mScarlet
DIC
crops
1
2
1
2
D
mito-mScarlet
DIC
E
mito-mScarlet
DIC

***Figure S5. Mitochondrial dynamics in oocyst development.*** *Live imaging of MitoRed oocysts on day 7 (A) day 10 (B) and day 13 (C) after mosquito infection. A) Oocyst at day 7 after infection with left image showing a maximum intensity projection of the mito-mScarlet signal. Right image shows a segmentation of the mito-mScarlet fluorescent signal by thresholding in Arivis software. Scale bar, 4 $\mu$m. C) two oocysts at day 13 after infection. Images on the right are crops of the mito-mScarlet signal of the image on the left, indicated by the dotted-line areas. Yellow arrowheads indicate mitochondrial organization centers (MOCs). D) Oocysts at day 13 showing beginning MOCs (yellow arrowheads). E) Oocyst at day 13 showing globular mitochondrial signal which could be a sign of unhealthy or dying parasites. B-E) Scale bars, 10 $\mu$m.*

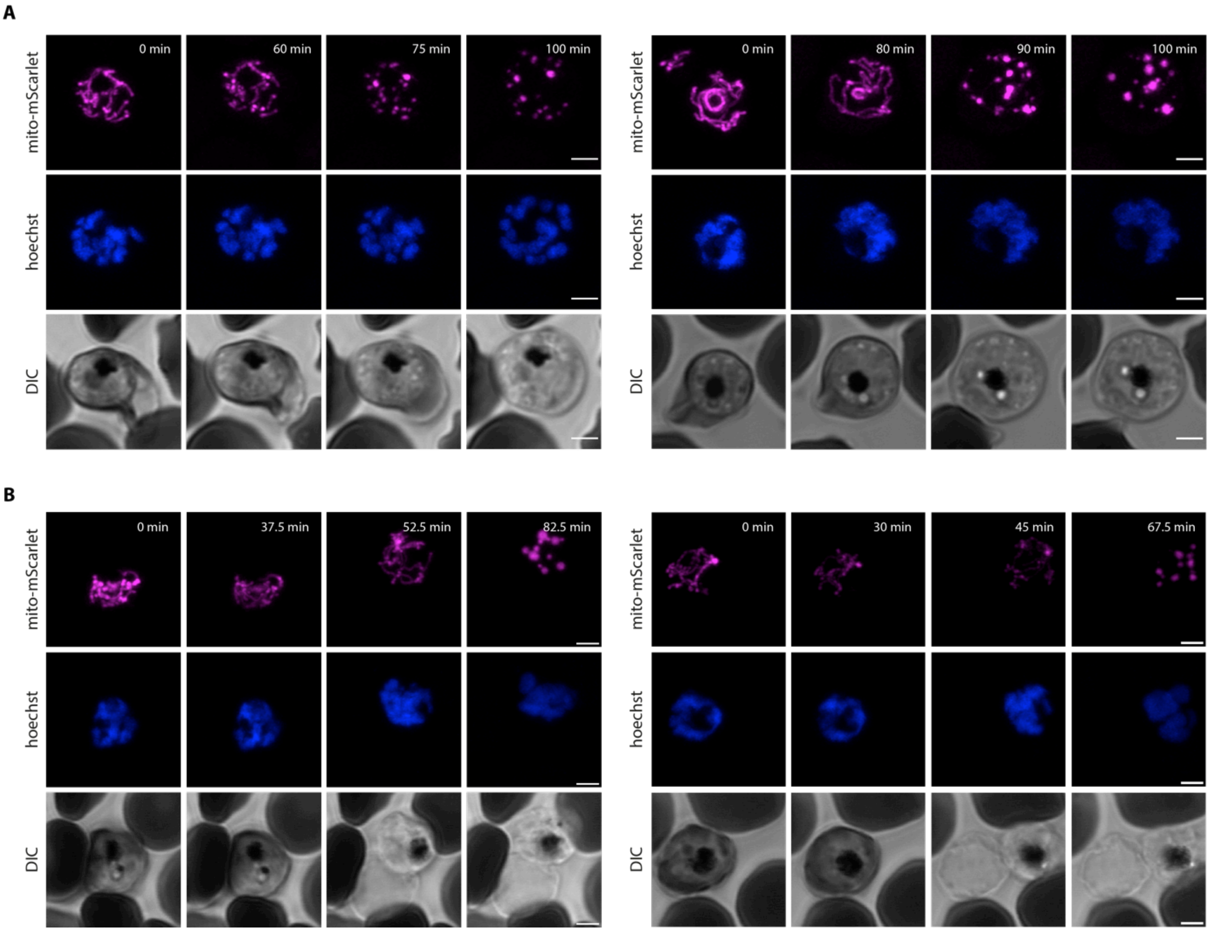


***Figure S6. Time-lapse imaging of MitoRed.*** *A) Live time-lapse imaging of MitoRed schizonts that show changes in parasite morphology. Mitochondria fall apart after approximately 75-90 minutes of imaging. B) Live time-lapse imaging of MitoRed schizonts that leave the RBC after 45-52.5 minutes of imaging. Parasites were stained with Hoechst to visualize DNA. All images are maximum intensity projections of Z-stacks taken with Airyscan confocal microscopy. Timestamps in the upper right corner represent the time points of the time-lapse experiment in minutes. Scale bars, 2 μm.*

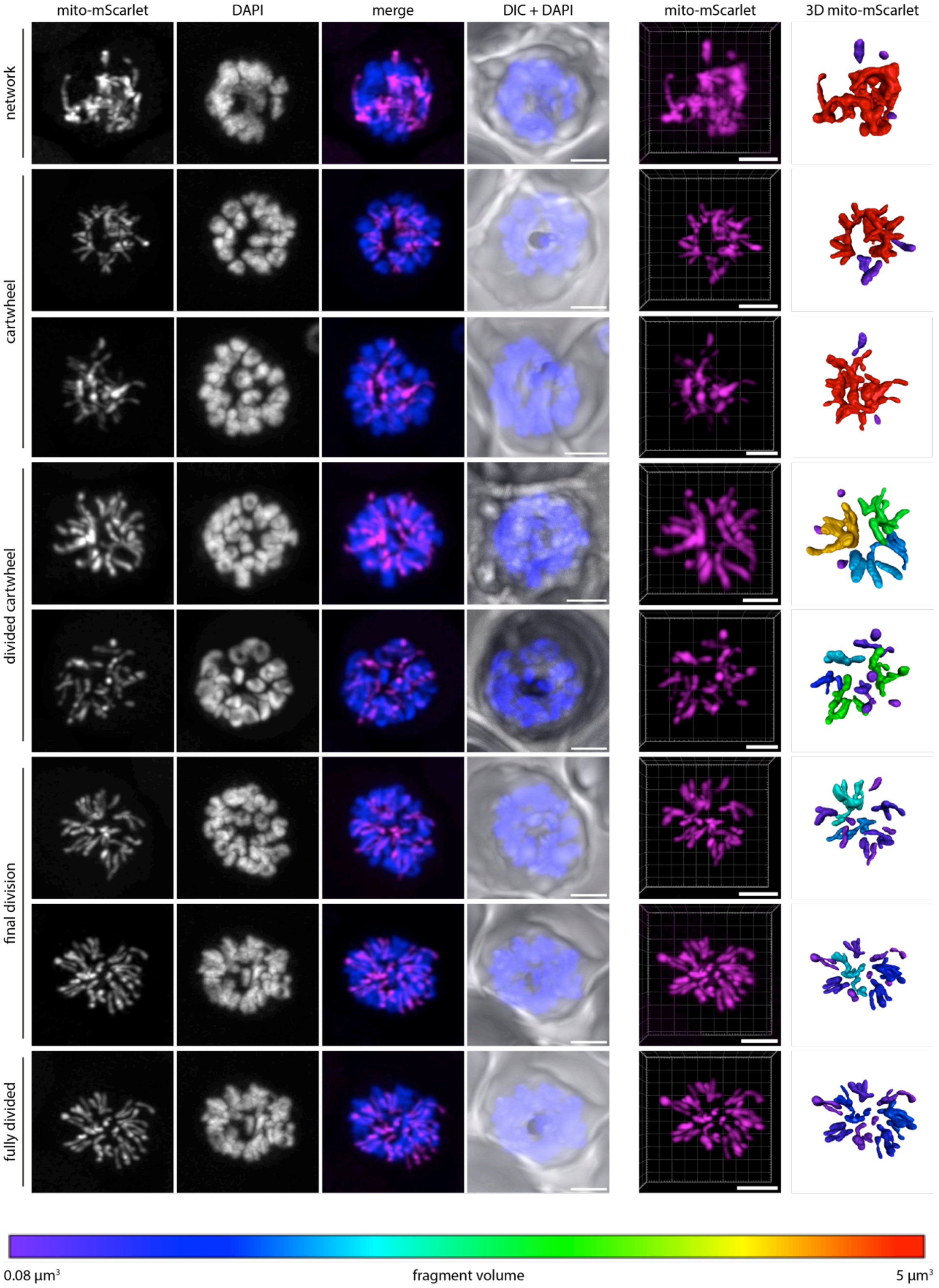


***Figure S7. Mitochondrial division stages in ABS schizogony.*** *Fluorescent imaging of MitoRed parasites in different mitochondrial division stages (described on the left). Images are representatives of the 17 parasites that were analyzed in second independent experiment. Mito-mScarlet signal is shown in magenta and DAPI (DNA) in blue. Images are maximum intensity projections of Z-stacks taken with an Airyscan confocal microscope. The fifth column shows the 3D image if the fluorescent mito-mScarlet signal, while the sixth column shows the 3D visualization of the segmented mitochondrial signal. The color of the mitochondrial fragment represents the size of this fragment, as is shown in the color bar at the bottom. Scale bars, 2 µm.*

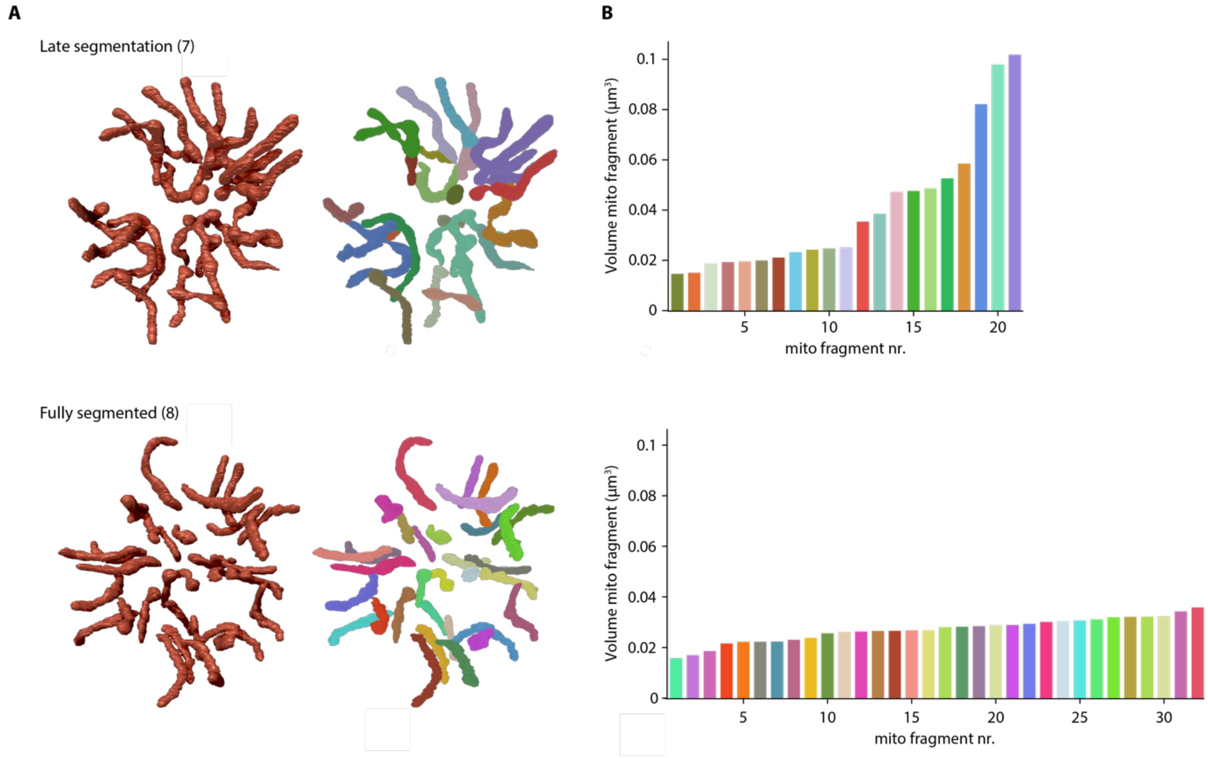


***Figure S8. Shape and volume of mitochondrial fragments during final stages of schizogony.*** *A) 3D rendering of mitochondria in a late and fully segmented schizont. The number between brackets indicates the parasite ID number (Table S3). In the right images, each mitochondrial fragment is depicted in an arbitrary color to distinguish separate and connected structures. B) Bar graphs indicating the mitochondrial fragment volumes in* $\mu m^3$ *and bar colors correspond to colors of the mitochondrial fragments in A.*

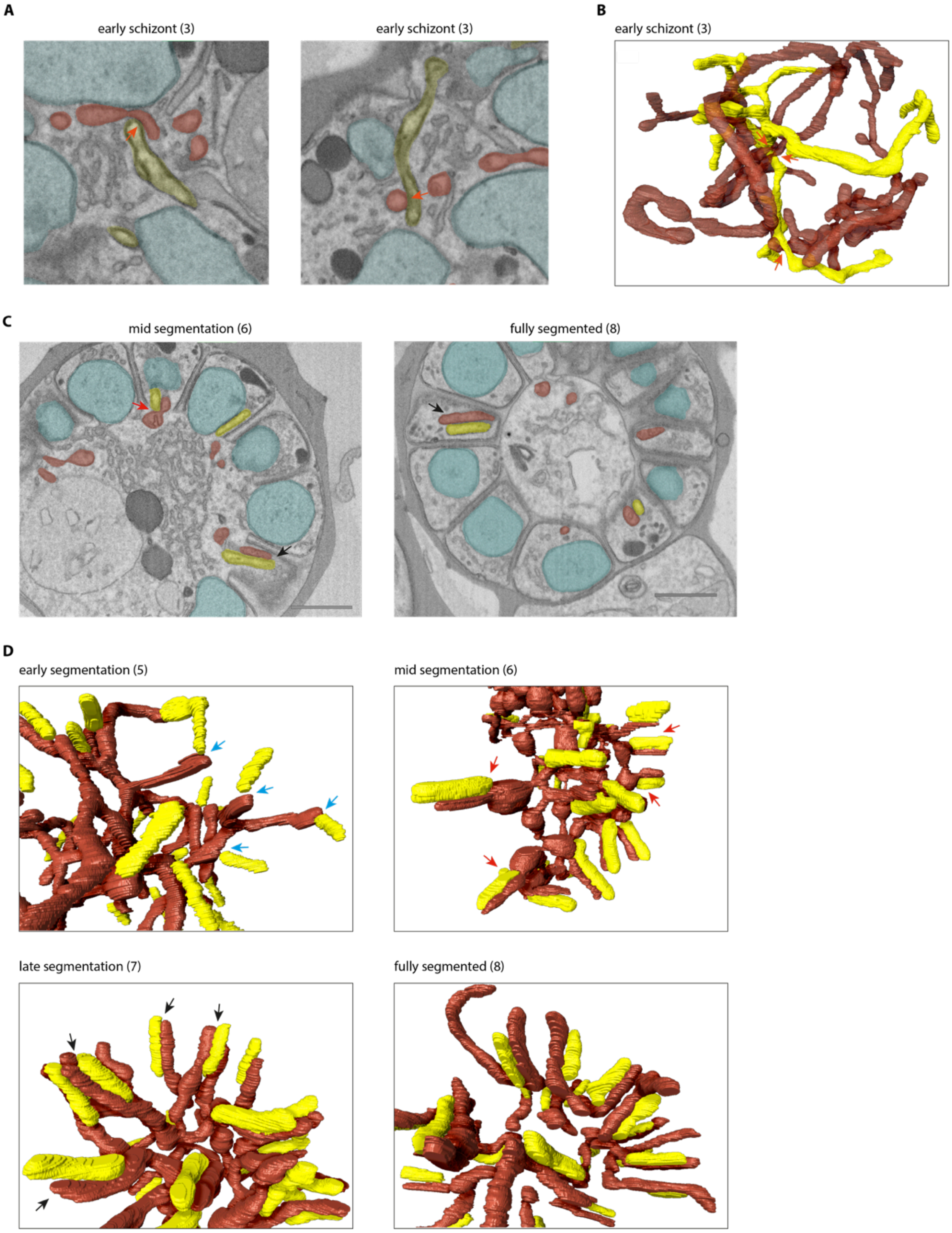


***Figure S9. Interaction between mitochondrion and apicoplast in different stages of schizogony.*** *A) Micrograph images of contact sites between the apicoplast (yellow) and mitochondrion (red) in early stage schizont, indicated by orange arrows. Nuclei are marked in teal. B) 3D rendering of mitochondrion (red, 50% opacity) and apicoplast (yellow), with contact sites indicated by orange arrows. C) Micrograph images of mid- and fully segmented schizonts showing contact sites where apicoplast and mitochondrion are aligned over the total apicoplast length (black arrows). Red arrow indicates where the basal end of the apicoplast is in contact with the bulbous part of the mitochondrion at the merozoite entrance. Scale bars, 1 μm. D) 3D rendering of apicoplast and mitochondrion in different stage schizonts. Blue arrows indicate where the basal end of an apicoplast interacts with the end of a mitochondrial branch. The number between brackets indicates schizont ID number (Table S3).*

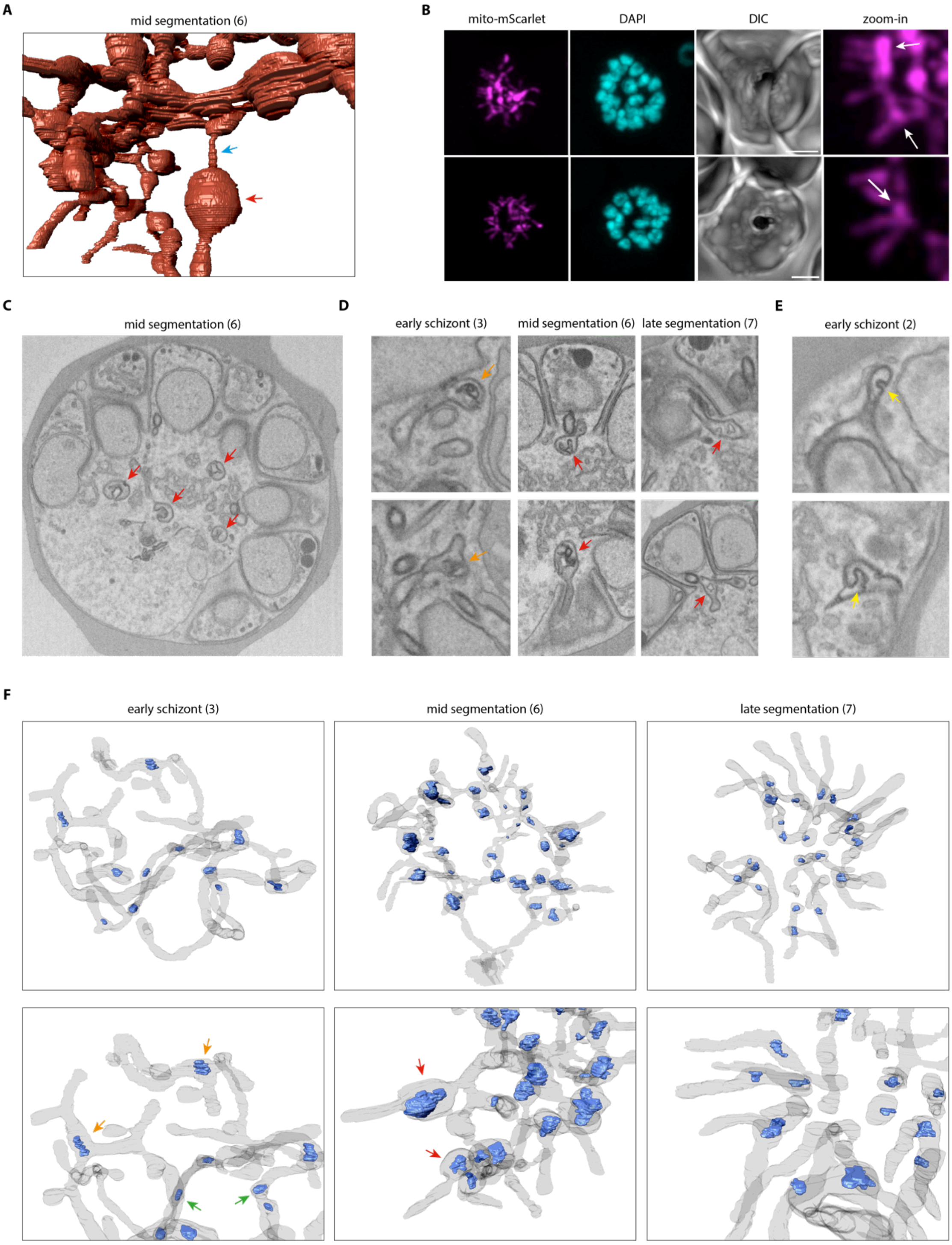


***Figure S10. Bulbous membrane invagination structures (bulins) in the mitochondrion.*** *A) 3D rendering of the mitochondrion in a mid-segmentation schizont with a thin (blue arrow) and thick (red arrow) part. B) Fluorescent microscopy of mito-mScarlet showing bulbous mitochondrial parts at the base of a mitochondrial branch. Scale bars, 2 μm. C) Micrograph image of mid-segmentation schizont showing four bulins. D) micrograph images of mitochondrial bulins in different schizont stages. E) micrograph images of apicoplast bulins (yellow arrows) in early stage schizont. F) 3D rendering of the mitochondrion (gray, 7% opacity) and membrane invaginations (blue). Red arrows indicate bulins at the base of a mitochondrial branch just outside the forming merozoite entrance. Orange arrows indicate membrane invaginations at a branching point in the mitochondrial network. Green arrows indicate membrane invaginations in the middle of a continuous mitochondrial branch. The number between brackets indicate schizont ID number (Table S3).*

**A** **B**

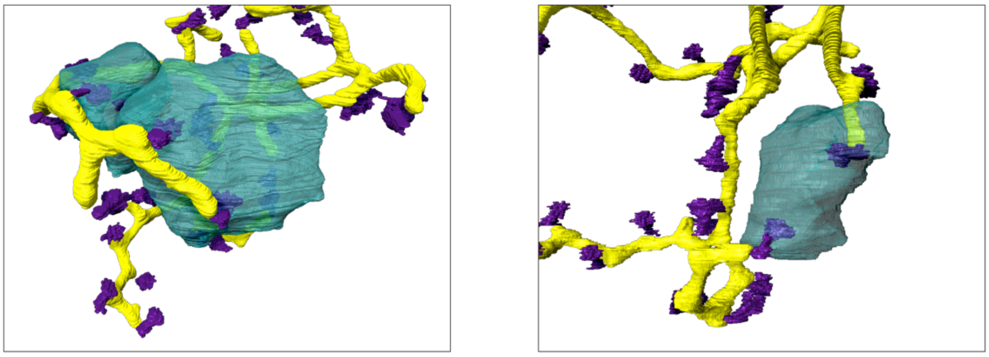

***Figure S11. Different interaction between apicoplast and centriolar plaques in an early schizont (3).*** *A) Two centriolar plaques (CPs, purple) from a single nucleus (teal) associating with one apicoplast branch (yellow). B) Two CPs from a single nucleus associating with two different apicoplast branches.*

### Movie legends

**Movie 1. Schizont 1.** Visualization of schizont 1 with parasite outline (gray), nuclei (cyan), apicoplast (yellow), mitochondrion (brown/red), centriolar plaques (purple).

**Movie 2. Schizont 2.** Visualization of schizont 2 with parasite outline (gray), nuclei (cyan), apicoplast (yellow), mitochondrion (brown/red), centriolar plaques (purple).

**Movie 3. Schizont 3.** Visualization of schizont 3 with parasite outline (gray), nuclei (cyan), apicoplast (yellow), mitochondrion (brown/red), centriolar plaques (purple).

**Movie 4. Schizont 4.** Visualization of schizont 4 with parasite outline (gray), nuclei (cyan), apicoplast (yellow), mitochondrion (brown/red), centriolar plaques (purple).

**Movie 5. Schizont 5.** Visualization of schizont 5 with parasite outline (gray), nuclei (cyan), apicoplast (yellow), mitochondrion (brown/red), centriolar plaques (purple).

**Movie 6. Schizont 6.** Visualization of schizont 6 with parasite outline (gray), nuclei (cyan), apicoplast (yellow), mitochondrion (brown/red), centriolar plaques (purple).

**Movie 7. Schizont 7.** Visualization of schizont 7 with parasite outline (gray), nuclei (cyan), apicoplast (yellow), mitochondrion (brown/red), centriolar plaques (purple).

**Movie 8. Schizont 8.** Visualization of schizont 8 with parasite outline (gray), nuclei (cyan), apicoplast (yellow), mitochondrion (brown/red), centriolar plaques (purple).

**Movie 9. Bulins during mid-segmentation.** Micrograph stacks of another mid-segmentation schizont showing merozoite-entrance bulins.

**Movie 10. Detailed bulins.** Micrograph stacks of schizont 6, showing two merozoite-entrance bulins, localizing at the basal end of the divided apicoplast.
